# Typical development of synaptic and neuronal properties can proceed without microglia in the cortex and thalamus

**DOI:** 10.1101/2024.09.06.611614

**Authors:** Mary O’Keeffe, Sam A Booker, Darren Walsh, Mosi Li, Chloe Henley, Laura Simões de Oliveira, Mingshan Liu, Xingran Wang, Maria Banqueri-Lopez, Katherine Ridley, Kosala Dissanyake, Cristina Martinez-Gonzales, Kirsty Craigie, Deepali Vasoya, Tom Leah, Xin He, David A Hume, Ian Duguid, Matthew F Nolan, Jing Qiu, David J. A. Wyllie, Owen R Dando, Alfredo Gonzales-Sulsar, Jian Gan, Clare Pridans, Peter C Kind, Giles E Hardingham

## Abstract

Brain-resident macrophages, microglia, have been proposed to play an active role in synaptic refinement and maturation, influencing plasticity and circuit-level connectivity. Here we show that a variety of neurodevelopmental processes previously attributed to microglia can proceed without them. Using a genetically modified mouse which lacks microglia (*Csf1r*^ΔFIRE/ΔFIRE^) we find that intrinsic properties, synapse number and synaptic maturation are largely normal in the hippocampal CA1 region and somatosensory cortex at stages where microglia have been implicated. Seizure susceptibility and hippocampal-prefrontal cortex coherence in awake behaving animals, processes disrupted in mice deficient in microglia-enriched genes, are also normal. Similarly, eye-specific segregation of inputs into the lateral geniculate nucleus proceeds normally in the absence of microglia. Single cell/nucleus transcriptomic analyses of neurons and astrocytes did not uncover any substantial perturbation due to microglial absence. Thus, the brain possesses remarkable adaptability to execute developmental synaptic refinement, maturation and connectivity in the absence of microglia.

## Introduction

The mammalian brain comprises complex networks of synaptically-connected neurons, supported by glia and the vasculature. During development there is a period of rapid synapse formation and elimination followed by refinement of synaptic connectivity and maturation of properties of surviving synapses, partly dependent on neuronal activity. Cell-autonomous mechanisms of neurons have been proposed to contribute these processes, including cytoskeletal rearrangements, pre/post-synaptic protein interactions, and membrane trafficking ^1^.

However, microglia have additionally been implicated in neuronal development, particularly in events taking place at the synapse ^2^. The majority of studies examining the role of microglia in brain development have focussed on three brain regions: hippocampal area CA1, the somatosensory cortex, and the lateral geniculate nucleus (LGN); examining the local and long-range connections of these brain areas ^3–13^. The contribution of microglia has been inferred from analyses of mice globally deficient in certain receptors highly (but not exclusively) expressed in microglia (Cx3cr1, Cr3, Trem2), employing complement component knockouts (C1q, C3), or by administering the *Csf1r* antagonist PLX5622 to kill microglia. While the results of these studies present a consistent role for microglia in developmental processes, further testing of this hypothesis is required as no model of microglial perturbation is without significant caveats. For example, the *Csf1r* antagonist PLX5622 has effects on non-microglial cells including circulating macrophages, T– and B-lymphocytes, and hematopoietic progenitors ^14^. Meanwhile, Cx3cr1 and the complement system can control metabolic and vascular properties, and aspects of brain development independently of microglial function ^15–22^.

Despite these caveats, a substantial body of work nevertheless indicates abnormal brain development when microglia function is altered ^3–13^. Here we have taken an alternative approach to addressing the role of microglia in brain development, by utilising *Csf1r*^ΔFIRE/ΔFIRE^ mice which are homozygous for the deletion of a promoter element (<u>F</u>ms-Intronic <u>R</u>egulatory <u>E</u>lement (FIRE) in the *Csf1r* gene ^23^. These mice lack microglia throughout life, while still possessing circulating monocytes and other CNS-resident (e.g. perivascular) macrophages ^23^. By focussing on developmental milestones and brain regions presumed disrupted by the presence of abnormal microglia, we provide the complementary approach of studying these processes in the absence of microglia.

## Results

### CA1 dendritic and synaptic properties in the absence of microglia

Prior to investigation of neuronal properties in *Csf1r*^ΔFIRE/ΔFIRE^ mice we confirmed an absence of microglia by RNA-seq, flow cytometry and immunohistochemistry (Extended Data Figs 1 and 2). RNA-sequencing (RNA-seq) analysis of *Csf1r*^ΔFIRE/ΔFIRE^ brain tissue at 14 and 42 postnatal days (P14 & P42) showed a depletion of microglia-specific genes including *C1q*, *Trem2* and *Cx3cr1* (Extended Data Fig. 1A, B). Immunohistochemical analysis of IBA1 expression in optically cleared brains confirmed microglial absence (Extended Data Fig. 2A), as did higher resolution imaging of the hippocampus, and neocortex (Extended Data Fig. 2B, C). Flow cytometry of dissociated P14 neocortex also confirmed the loss of CD11b^+^/CD45^low^ microglia (Extended Data Fig. 1C, D), but retention of (far less numerous) CD11b^+^/CD45^high^ macrophages (Extended Data Fig. 1C) and other glia (Extended Data Fig. 1E, F). *Csf1r*^ΔFIRE/ΔFIRE^ mice have normal body and brain weight ^23^, avoiding confounds of other microglia-deficient models, such as premature death, bone abnormalities, and altered CNS macrophages and monocytes ^14,24,25^. *Csf1r*^ΔFIRE/ΔFIRE^ pups appeared healthy and developed typical innate motor skills, as tested by righting reflex and negative geotaxis tasks (Extended Data Fig. 3A-C).

As shown above, the fractalkine receptor *Cx3cr1* is an established microglial signature gene, and is almost entirely absent in *Csf1r*^ΔFIRE/ΔFIRE^ brains (Extended Data Fig. 1A, B). Mice deficient in CX3CR1 were reported to show transient reductions of microglia P8 to P28 in *stratum radiatum* of hippocampal area CA1, coinciding with elevated synapse number at P15 as assessed by physical (spine density) and functional analysis (mEPSC frequency and amplitude) ^3^. CA1 pyramidal cells (PCs) of *Csf1r*^ΔFIRE/ΔFIRE^ mice were morphologically similar to wild-type, with dendritic arborization indistinguishable between the two genotypes at P14 (Extended Data Fig. 4A-D) and P28 (Extended Data Fig. 4E-H). We also found no difference in the density of putative dendritic spines on basal, oblique or tuft dendrites of CA1 PCs at P14 or P28 (Fig. 1A, B). We next performed stimulated emission depletion (STED) microscopy to visualise the structure of dendritic spines in *Csf1r*^ΔFIRE/ΔFIRE^ mice beyond the diffraction limit of light. At P14 in CA1 PCs, we found no genotype-dependent differences in spine overall length, head width, or neck length (Fig. 1C, D), and we confirmed that spine density was not altered (Fig. 1E). Finally, we confirmed that functional synapses were not impaired, by measuring miniature EPSC frequency or amplitude in CA1 PCs. We found no difference in the quantal strength (mEPSC amplitude) or number (mEPSC frequency) between *Csf1r*^ΔFIRE/ΔFIRE^ and *Csf1r*^+/+^ mice at either P14 or P28 (Fig. 1F-H). These data show that dendritic spine density and morphology, and also synaptic function, was unaffected by an absence of microglia at developmental stages where the differences have been previously reported ^3^.

**Figure 1.**
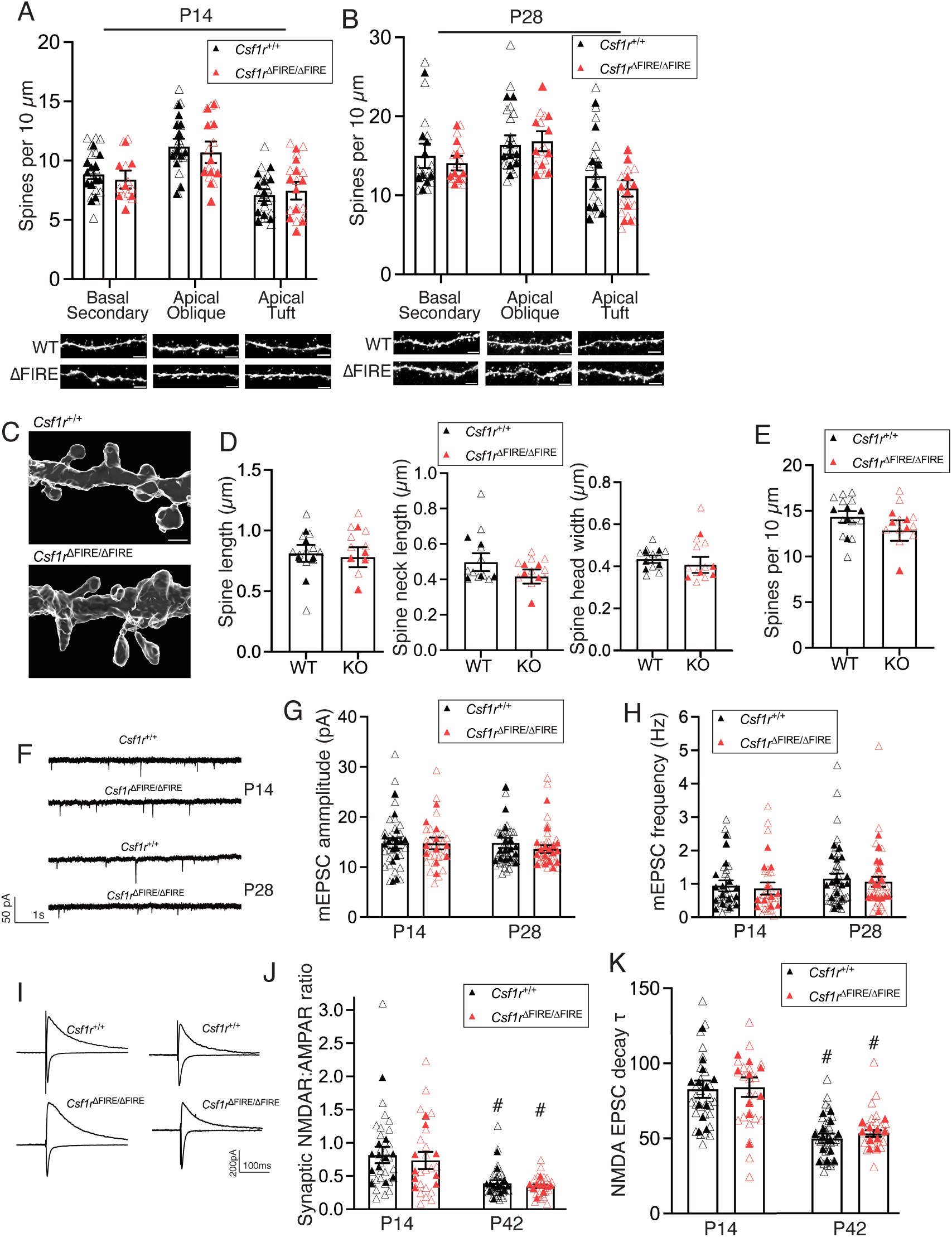
Absence of brain microglia does not alter hippocampal synaptic density or properties. All data are shown as mean ± SEM and all statistical tests are two-sided. **A**) CA1 spine density (P14). Basal secondary: (F_(1,17)_=0.825, t_(17)_=0.908, p=0.377, nested Student’s t-test, n=12 *Csf1r^+/+^* (5m/7f) and n=7 *Csf1r*^ΔFIRE/ΔFIRE^ mice (5m, 2f). Apical oblique: (F_(1,20)_=0.257, t_(20)_ 0.507, p=0.617, nested Student’s t-test, n=12 *Csf1r*^+/+^ (5m, 7f) and n=10 *Csf1r*^ΔFIRE/ΔFIRE^ mice (6m/4f). Apical tuft: (F_(1,19)_=0.356, t_(19)_ 0.597, p=0.558, nested Student’s t-test, n=11 *Csf1r*^+/+^ (5m/6f) and n=10 *Csf1r*^ΔFIRE/ΔFIRE^ mice (6m, 4f). Scale bars: 3 µm. **B**) Spine density at P28. Basal secondary: (F_(1,15)_=0.260, t_(15)_=0.510, p=0.618, nested Student’s t-test, p>0.999 Mann Whitney test, n=9 *Csf1r*^+/+^ (3m, 6f) and 8 *Csf1r*^ΔFIRE/ΔFIRE^ mice (4M, 4F). Apical oblique: (F_(1,16)_=0.001, t_(16)_=0.036, p=0.972, nested Student’s t-test, 0.667 Mann Whitney test, n=9 *Csf1r*^+/+^ (3m, 6f) and 9 *Csf1r*^ΔFIRE/ΔFIRE^ mice (5m, 4f). Apical tuft dendrites: (F_(1,16)_=0.872, t_(16)_=0.934, p=0.364, nested Student’s t-test, 0.667 Mann Whitney test, n=9 *Csf1r*^+/+^ (3m, 6f) and 9 *Csf1r*^ΔFIRE/ΔFIRE^ mice (5m, 4f). Here and throughout the manuscript filled symbols represent independent biological replicates (animals) and open symbols represent technical replicates (multiple cells studied per animal). **C)** Example STED images of CA1 apical oblique dendrites (P28). Scale bar: 700 nm. **D)** STED microscopy of sections of CA1 apical oblique dendrites (P28) and analysis of spine length (left graph: t_(8)_=0.297, p=0.774), spine neck length (middle graph, t_(8)_=1.26, p=0.244) and spine head width (right graph, t_(8)_=0.646, p=0.537); n=5 *Csf1r*^+/+^ (2m/3f) and 5 *Csf1r*^ΔFIRE/ΔFIRE^ mice (2m/3f). **E)** Spine density of CA1 apical oblique dendrites derived from STED images: t_(8)_=1.16, p=0.280 (unpaired t-test); n=5 *Csf1r*^+/+^ (2m/3f) and 5 *Csf1r*^ΔFIRE/ΔFIRE^ mice (2m/3f). **F)** mEPSCs recorded at P14 (upper) and P28 (lower). **G, H**) Amplitude (G) of mEPSCs at P14 (F_(1,25)_=0.345, t_(25)_ 0.588, p=0.562, nested Student’s t-test) and P28 (F_(1,31)_=0.363, t_(31)_=0.603, p=0.551, nested Student’s t-test; p=0.345 Mann Whitney test). mEPSC frequency (H) at P14 (F_(1,25)_=0.0404, t_(25)_=0.2010, p=0.842, nested Student’s t-test) and P28 (F_(1,31)_=0.157, t_(31)_=0.397, p=0.694 nested Student’s t-test, p=0.625 Mann Whitney test). P14: n= 16 *Csf1r*^+/+^mice (7m, 8f, 1u/n), n= 11 *Csf1r*^ΔFIRE/ΔFIRE^ mice (4M, 4F, 3 u/n); P28: n= 16 *Csf1r*^+/+^mice (7m, fF), n=17 *Csf1r*^ΔFIRE/ΔFIRE^ mice (7m, 10f) **I**) Example EPSC traces. AMPAR and NMDAR-mediated EPSCs were recorded at –70 mV and +40 mV respectively. **J)** Quantification of the NMDAR/AMPAR ratios at P14 (F_(1,19)_=0.110, t_(19)_=0.332, p=0.743, nested Student’s t-test) and P42 (F_(1,21)_=0.623, t_(21)_=0.789, p=0.439, nested Student’s t-test, p=0.879 Mann Whitney test). P14: n=12 *Csf1r*^+/+^ (6m/6f) and 9 *Csf1r*^ΔFIRE/ΔFIRE^ mice (6m/3f); P42: n=13 *Csf1r*^+/+^ (6m/7f) and 10 *Csf1r*^ΔFIRE/ΔFIRE^ mice (5m/5f). # p<0.0001 (Main age effect, 2-way ANOVA on the averages of each of the animals), Šídák’s multiple comparisons test: p=0.0022 (*Csf1r*^+/+^), 0.0158 (*Csf1r*^ΔFIRE/ΔFIRE^). **K)** Quantification of the decay constant of NMDAR-mediated EPSCs at P14 (F_(1,19)_=0.0008, t_(19)_=0.027, p=0.978, nested Student’s t-test) and P42 (F_(1,21)_=0.737, t_(21)_=0.859, p=0.400, nested Student’s t-test). P14: n= 12 *Csf1r*^+/+^ (6m/6f) and 9 *Csf1r*^ΔFIRE/ΔFIRE^ mice (5m/6f); P42: n= 13 *Csf1r*^+/+^ (6m/7f) and 10 *Csf1r*^ΔFIRE/ΔFIRE^ mice (5m/5f). # p<0.0001 (Main age effect, 2-way ANOVA performed on the averages of each of the animals), Šídák’s multiple comparisons test: p<0.0001 (*Csf1r*^+/+^), 0.0002 (*Csf1r*^ΔFIRE/ΔFIRE^).

In addition to a presumed role in controlling CA1 synapse number in early development, microglia have been implicated in regulating maturation of synapses – in terms of neurotransmitter receptor content and presynaptic function ^12^. In putative CA3-CA1 connections measured by stimulating the *stratum radiatum* in brain slices from *Csf1r*^ΔFIRE/ΔFIRE^ mice, we observed normal development of NMDAR:AMPAR ratio from P14 to P42 (Fig. 1I, J). We also confirmed that developmental regulation of NMDA receptor-mediated EPSCs kinetics, a measure of NMDAR subunit composition, finding the temporal reduction in decay time-constant to be similar in recordings of CA1 PCs from *Csf1r*^ΔFIRE/ΔFIRE^ and *Csf1r*^+/+^ mice. (Fig. 1K). Consistent with similar synaptic properties in the absence of microglia, we also observed normal synaptic plasticity: induction of long-term potentiation (LTP) and depression (LTD) at Schaffer-collateral synapses in *Csf1r*^ΔFIRE/ΔFIRE^ mice was similar to that of *Csf1r*^+/+^ mice P14 (Extended Data Fig. 5A, B). In addition to basal synaptic properties of neurons, how neurons intrinsically respond to synaptic inputs is a critical determinant of brain circuit development. We therefore measured the intrinsic physiological properties of *Csf1r*^ΔFIRE/ΔFIRE^ CA1 pyramidal neurons at the age (P42) studied above. We found intrinsic properties in *Csf1r*^ΔFIRE/ΔFIRE^ neurons (frequency-current curve, resting membrane potential, input resistance, rheobase, action potential threshold, paired-pulse ratio) to be indistinguishable from wild-type (Fig. 2A-F). Thus, microglia are not essential for several aspects of hippocampal CA1 synaptic and cellular development for which they have been proposed to play a role.

**Figure 2.**
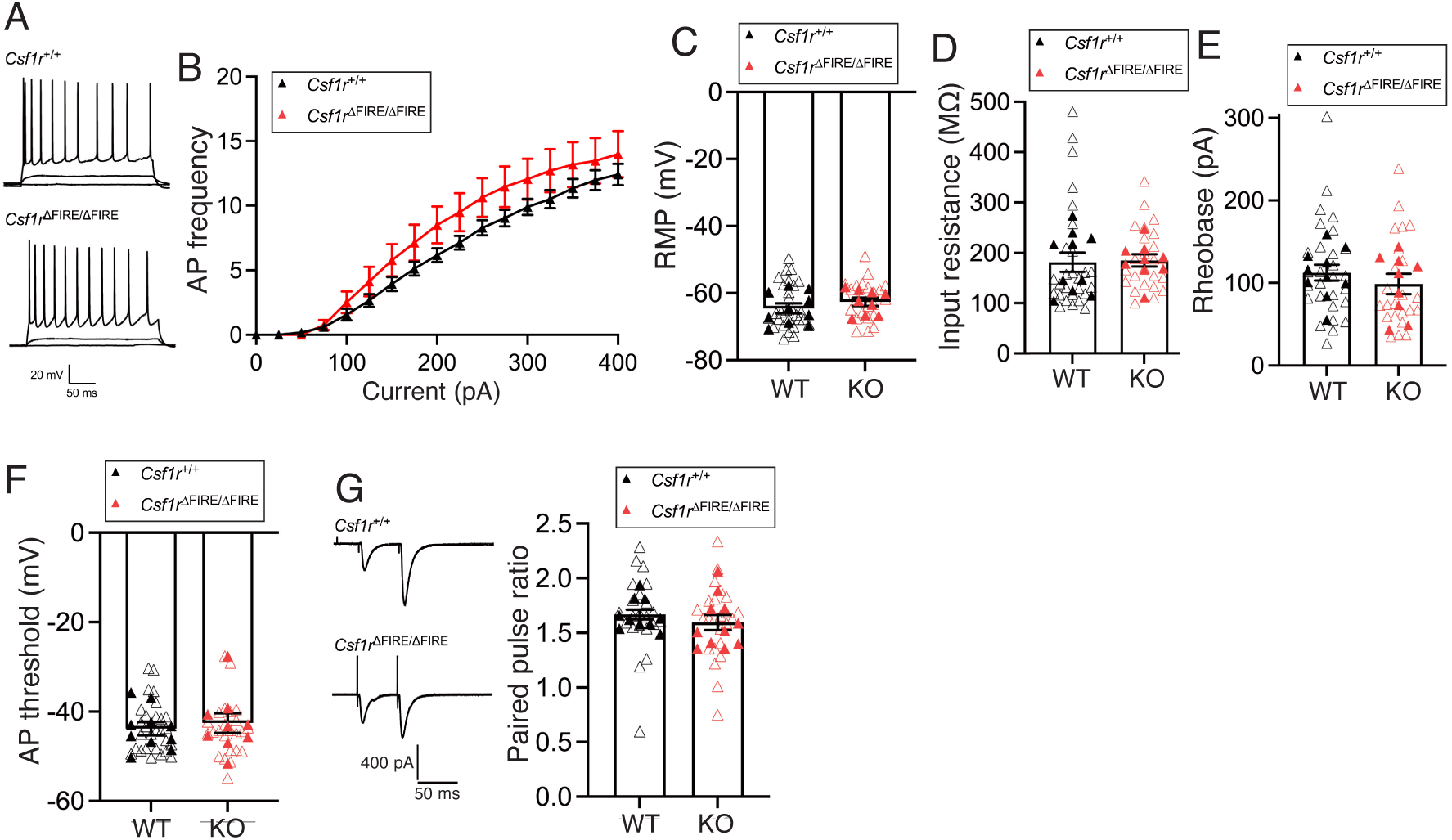
No change in intrinsic neuronal excitability or short term plasticity in CA1 of the hippocampus of mice lacking microglia. All data are shown as mean ± SEM and all statistical tests are two-sided. **A**) Representative traces of the membrane voltage of CA1 pyramidal cells at P42 in *Csf1r*^+/+^ (upper) and *Csf1r*^ΔFIRE/ΔFIRE^ (lower) mice in response to hyper-to depolarising current steps (−100 to +400 pA, 25 pA steps, 500 ms duration). **B)** AP frequency/current relationship (F/I curve), (F_(1,17)_=1.60 p=0.223, repeated measures 2-way ANOVA [genotype effect], n=10 *Csf1r*^+/+^ mice (7m/3f) and n=9 *Csf1r*^ΔFIRE/ΔFIRE^ mice (5m/4f). Absence of microglia during early development had no discernible effect on CA1 pyramidal resting membrane potential (**C**, F_(1,17)_=1.09, t_(17)_=1.04, p=0.311, nested Student’s t-test), input resistance (**D**, F_(1,17)_=0.032, t_(17)_=0.178, p=0.861, nested Student’s t-test), rheobase (**E**, F_(1,17)_=1.09, t_(17)_=1.05, p=0.310, nested Student’s t-test), or action potential (AP) threshold (**F**, F_(1,17)_=0.045, t_(17)_=0.212, p=0.834, nested Student’s t-test. For D-F, n=10 (7m/3f) *Csf1r*^+/+^ and 9 (5m/4f) *Csf1r*^ΔFIRE/ΔFIRE^ mice. **G**) Delivery of paired-pulse electrical stimulation (2x stimuli, 50 ms interval) to the Schaffer-Collaterals resulted in facilitating EPSCs, which did not differ between genotypes. F_(1,19)_=0.413, t_(19)_=0.413 p=0.528, nested Student’s t-test (n=10 *Csf1r*^+/+^ mice (6m/4f) and n=11 *Csf1r*^ΔFIRE/ΔFIRE^ mice (5m/6f).

### Refinement of eye-specific inputs in the LGN without microglia

Besides Cx3cr1 signaling, several studies propose that microglia-mediated synapse removal depends on mechanisms involving components of the complement system, including C1q (an initiator of the classical complement cascade) and C3 (a target for phagocytosis by CR3-expressing cells such as microglia). This system has been implicated in the axonal refinement of retinal ganglion cells in the lateral geniculate nucleus (LGN), where inputs from both eyes overlap in early development, but later segregate into non-overlapping eye-specific zones ^26–29^. Studies have reported that this segregation proceeds normally in *Cx3cr1*^-/-^ mice ^30^, but is impaired in mice deficient in CR3, C3 or C1qa ^2,7,8^. *Csf1r*^ΔFIRE/ΔFIRE^ mice express >95% lower *C1qa/b/c* mRNA in the brain (Extended Data Fig. 1A, B). Consistent with previous studies ^26–29^ we observe that inputs into the LGN from both eyes overlap early in development (P4), and by P10 have segregated into eye-specific fields (Fig. 3A, B, E). However, we found that developmental eye-specific segregation in the LGN is normal in *Csf1r*^ΔFIRE/ΔFIRE^ mice lacking microglia (Fig. 3C, D, E). Thus, there is no noticeable requirement for microglia for developmental axon terminal refinement in the LGN.

**Figure 3.**
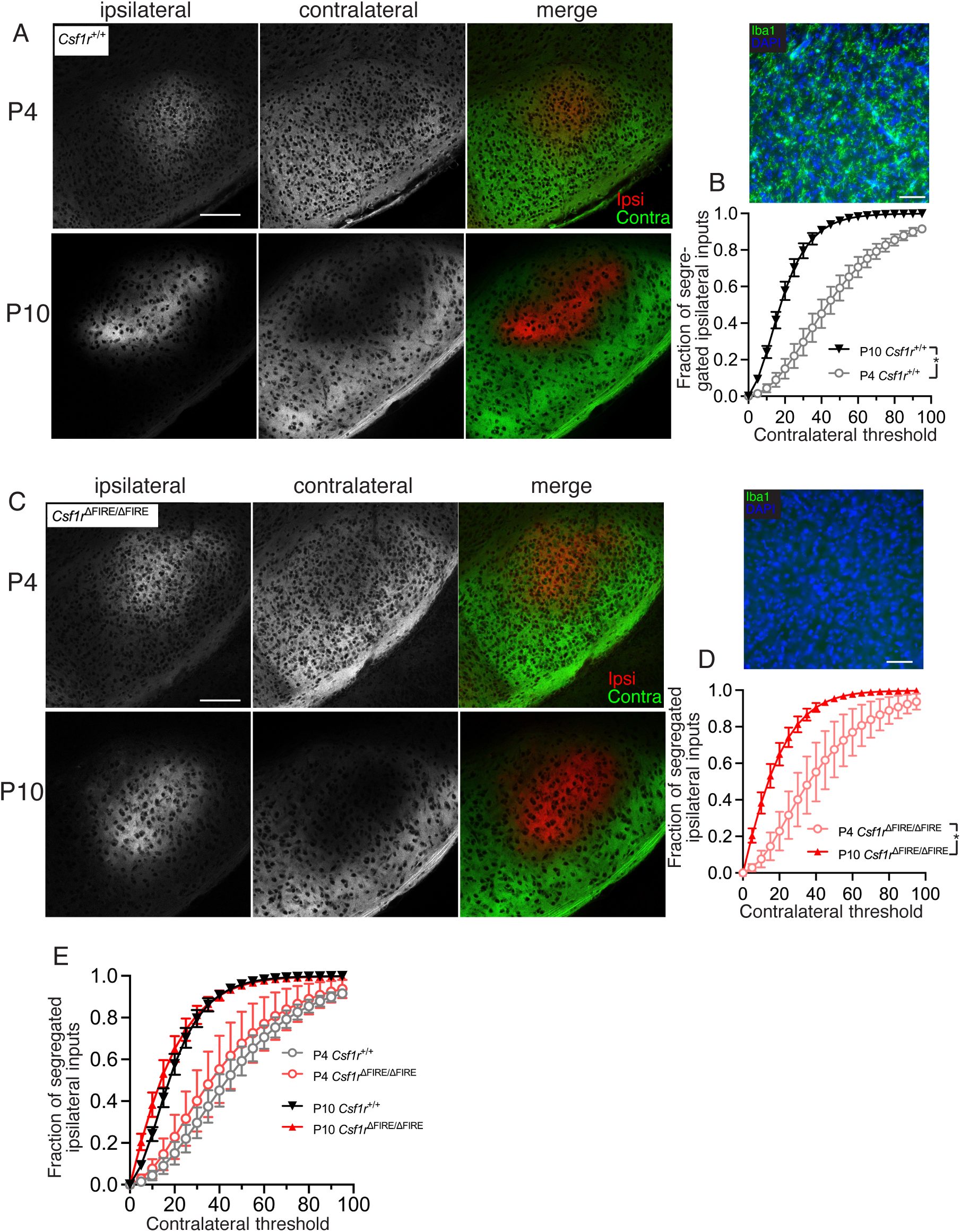
Retinal inputs into the LGN segregate typically in the absence of brain microglia. All data are shown as mean ± SEM and all statistical tests are two-sided. **A**) Example confocal images of the dorsal lateral geniculate nucleus (dLGN) of P4 and P10 *Csf1r*^+/+^ mice following anterograde labelling of inputs from the ipsilateral (DiI, red) and contralateral (DiO, green) retinas. Scale bar: 100 µm. **B**) Quantification of dLGN segregation by measuring the fraction of segregated inputs at P4 (open circles) and P10 (closed triangles) in *Csf1r*^+/+^ mice (F_(1,12)_=46.97, p=1.8E-03, 2-way ANOVA [age effect); n=5 mice (P4, 3m/2f), 9 mice (P10, 3m/6f). Upper shows example image of DAPI/IBA1 staining (scale bar: 50 µm). **C**) Example images of dLGN segregation in *Csf1r*^ΔFIRE/ΔFIRE^ mice according to the same scheme as **A**. Scale bar: 100 µm. **D**) dLGN segregation measured at P4 (open circles) and P10 (closed triangles) in *Csf1r*^ΔFIRE/ΔFIRE^ mice (F_(1,14)_=14.07, p=0.002, 2-way ANOVA [age effect]), n=4 (P4, 2m/2f) and 12 (P10, 6m/6f). Upper shows example image of DAPI/IBA1 staining (scale bar: 50 µm). **E**) Comparison of the dLGN input segregation at P10 (F_(1,19)_=0.686, p=0.418, 2-way ANOVA [genotype]) and P4 (F_(1,7)_=0.351, p=0.572, 2-way ANOVA [genotype]) displayed no difference between genotypes (for n numbers see B and D).

### The barrel cortex of Csf1r^ΔFIRE/ΔFIRE^ mice develops normally

In addition to the LGN, microglia have also been implicated in refinement of synapses in the barrel-field of the primary somatosensory (S1) cortex ^4,10^. The barrel field is known to undergo stereotypical patterning in layer 4 (L4) between P4 and P10 involving axonal and dendritic refinement. *Csf1r*^ΔFIRE/ΔFIRE^ mice displayed normal anatomical organisation of the barrel field compared to *Csf1r*^+/+^ littermates, both in terms of overall barrel field size, but also barrel organisation, and synaptic labelling (Fig. 4A-D, Extended Data Fig. 6A-C). To next determine, if basal synaptic transmission was altered in L4 of S1, as has been suggested following microglia depletion by administration of *CSF1R* inhibitor PLX5622 ^10^. We found that P14 *Csf1r*^ΔFIRE/ΔFIRE^ and *Csf1r*^+/+^ L4 S1 neurons had similar spontaneous EPSC and IPSC amplitude and frequencies (Fig. 4E-J), leading us to conclude that excitatory and inhibitory synaptic development is not strongly affected by the constitutive absence of microglia.

**Figure 4.**
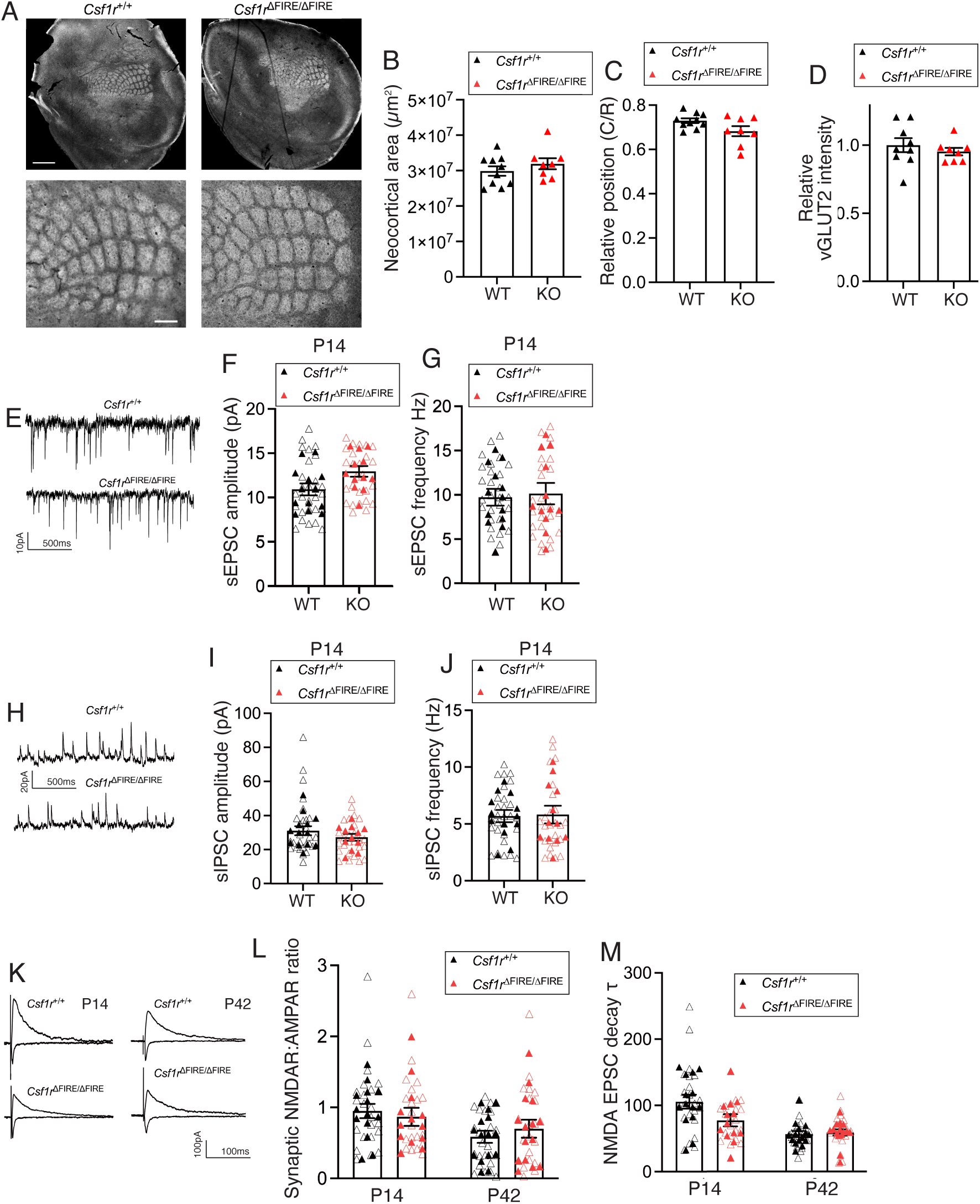
Assessment of synaptic properties in the somatosensory cortex development in the absence of microglia. All data are shown as mean ± SEM and all statistical tests are two-sided. **A**) Example VGLUT2 immunofluorescent images of the somatosensory barrel cortex from *Csf1r*^+/+^ (left) and *Csf1r*^ΔFIRE/ΔFIRE^ (right) mice. Scale bars: 1 mm (upper) and 200 µm (lower). **B)** Quantification of neocortical area occupied by the barrel-field (t_(16)_=1.01, p=0.329, Student’s 2-tailed t-test), n=10 *Csf1r*^+/+^ and 8 *Csf1r*^ΔFIRE/ΔFIRE^ mice. **C**) Caudal/rostral relative position of the barrel field (t_(16)_=2.01, p=0.062, Student’s 2-tailed t-test), n=10 *Csf1r*^+/+^ and 8 *Csf1r*^ΔFIRE/ΔFIRE^ mice. **D)** VGLUT2 staining intensity within the barrel cortex: t_(15)_=0.78, p=0.448, Student’s t-test, n=9 *Csf1r*^+/+^ and 8 *Csf1r*^ΔFIRE/ΔFIRE^ mice. **E**) Example spontaneous EPSC recordings from P14 L4 stellate cells at –70 mV. **F, G)** No difference in sEPSC amplitude (**F**, F_(1,23)_=3.36, t_(23)_=1.83, p=0.081, nested Student’s t-test) or frequency (**G**, F_(1,23)_=0.029, t_(23)_=0.171, p= 0.866, nested Student’s t-test), n=13 *Csf1r*^+/+^ mice (8m/5f) and 12 *Csf1r*^ΔFIRE/ΔFIRE^ mice (6m/6f). **H**) Example spontaneous inhibitory postsynaptic potential (sIPSC) recordings from P14 L4 stellate cells. **I, J)** No difference in sIPSC amplitude (**I**, F_(1,23)_=2.05, t_(23)_=1.43, p=0.166, nested Student’s t-test) or frequency (**J**, F_(1,23)_=0.003, t_(23)_=0.050, p=0.961, nested Student’s t-test); n= 13 *Csf1r*^+/+^ mice (8m/5f) and 12 *Csf1r*^ΔFIRE/ΔFIRE^ mice (6m/6f). **K**) (Left) Example traces of EPSCs recorded in L4 stellate cells. **L)** AMPAR and NMDAR mediated EPSCs were measured following stimulation of thalamo-cortical afferents. Quantification of the NMDAR:AMPAR EPSC ratios confirmed no genotype difference at P14 (F_(1,25)_=0.330, t_(25)_=0.574, p=0.571, nested Student’s t-test) or P42 (F_(1,28)_=0.809, t_(28)_=0.899, p=0.376, nested Student’s t-test). P14: n=14 *Csf1r*^+/+^ mice (7m/7f) and 13 *Csf1r*^ΔFIRE/ΔFIRE^ mice (8m/5f); P42: n=15 *Csf1r*^+/+^ (8m/7f) and 15 *Csf1r*^ΔFIRE/ΔFIRE^ mice (6m/9f). **M**) Measurement of NMDAR-mediated EPSC decay time-constant (τ) revealed a small difference at P14 (F_(1,25)_=6.69, t_(25)_=2.59, p=0.016, nested Student’s t-test; n=14 *Csf1r*^+/+^ (7m/7f) and 13 *Csf1r*^ΔFIRE/ΔFIRE^ mice (8m/5f). No difference was observed at P42 (F_(1,28)_=0.410, t_(28)_=0.641, p=0.527, nested Student’s t-test; n=15 *Csf1r*^+/+^ (8m/7f) and 15 *Csf1r*^ΔFIRE/ΔFIRE^ mice (6m/9f).

We next examined the synaptic AMPAR:NMDAR EPSC ratio and functional NMDAR kinetics in L4 S1, since these properties were reported to be altered in *Cx3cr1^−/−^* mice ^4^. We observed that the AMPAR:NMDAR ratio of S1 L4 neurons was similar in *Csf1r*^ΔFIRE/ΔFIRE^ and *Csf1r*^+/+^ at both P14 and P42 (Fig. 4K, L). Interestingly, the weighted time constant of NMDAR-mediated EPSCs, which reflects the subunit composition of NMDARs, was found to be slightly lower in *Csf1r*^ΔFIRE/ΔFIRE^ neurons at P14 than wild-type, although this difference was no longer apparent at P42 (Fig. 4M). Furthermore, the intrinsic physiological properties of L4 neurons (frequency-current curve, resting membrane potential, input resistance, rheobase, action potential threshold) were indistinguishable from those in *Csf1r*+/+ mice, as was paired-pulse ratio (Fig. 5A-H). Overall, our data demonstrate that the absence of microglia does not compromise normal anatomical or electrophysiological development of the barrel cortex, with the exception of a precociously low decay time constant in NMDA receptor-mediated synaptic responses at P14.

**Figure 5:**
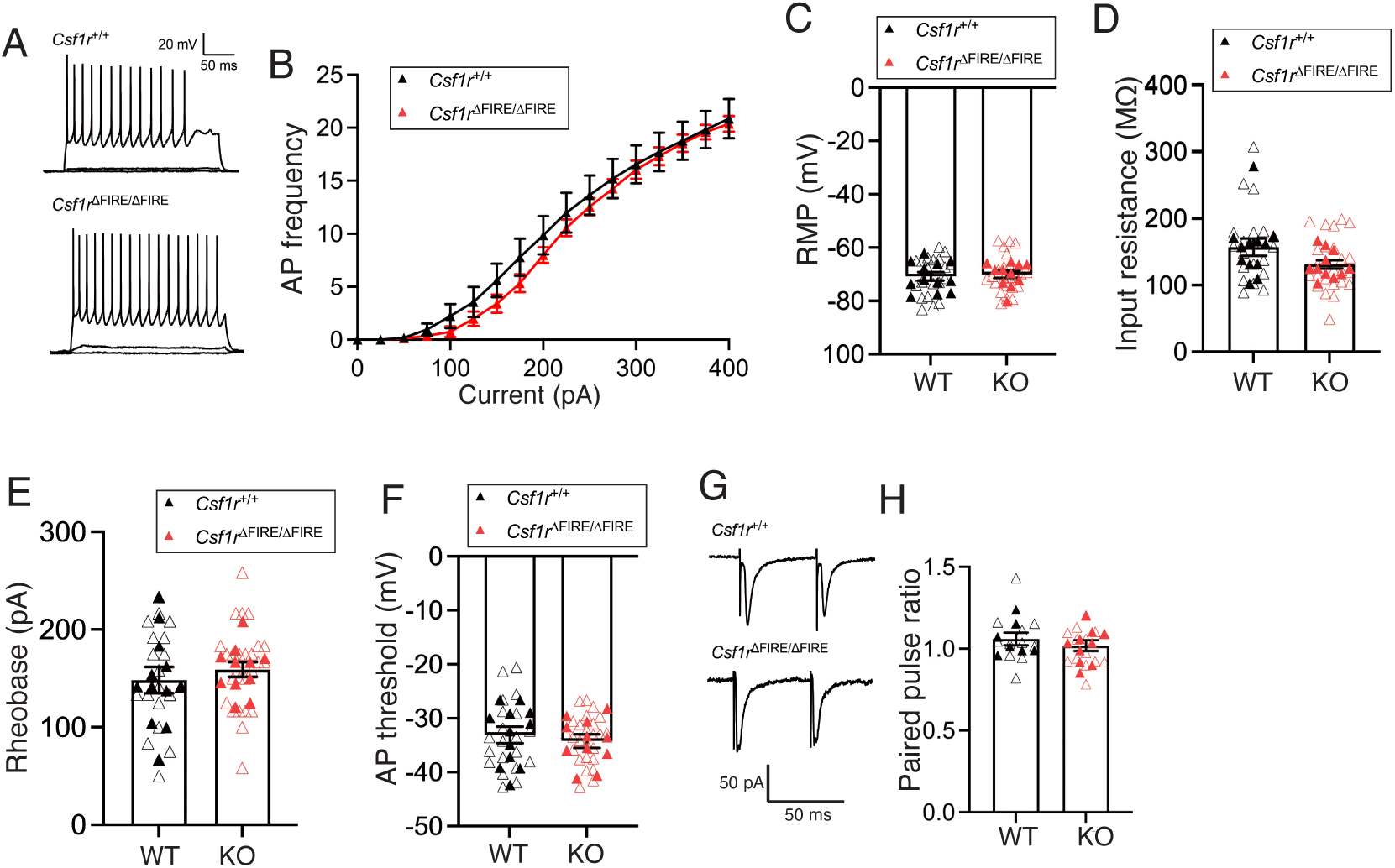
No change in intrinsic neuronal excitability or short term plasticity in the somatosensory cortex of mice lacking microglia. All data are shown as mean ± SEM and all statistical tests are two­sided. **A**) Representative traces of the membrane voltage of L4 stellate cells at P42 in *Csf1r*^+/+^ (upper) and *Csf1r*^ΔFIRE/ΔFIRE^ (lower) mice in response to hyper-to depolarising current steps (−100 to +400 pA, 25 pA steps, 500 ms duration). **B)** AP frequency/current relationship (F/I curve), (F_(1,21)_=0.401 p=0.533, repeated measures 2-way ANOVA [genotype effect], n=12 *Csf1r*^+/+^ mice (9m/3f) and n=11 *Csf1r*^ΔFIRE/ΔFIRE^ mice (7m/4f). We observed no change in resting membrane potential (**C**, RMP, F_(1,21)_=0.216, t_(21)_=0.465, p=0.647, nested Student’s t-test), input resistance (**D**, F_(1,21)_=3.51, t_(21)_=1.87, p=0.075, nested Student’s t-test), rheobase (**E**, F_(1,21)_=0.766, t_(21)_=0.875, p=0.391, nested Student’s t-test), AP threshold (**F**, (F_(1,21)_=0.206, t_(21)_=0.454, p=0.655, nested Student’s t-test). For C-F n=12 *Csf1r*^+/+^ (9m/3f) and n=11 *Csf1r*^ΔFIRE/ΔFIRE^ (7m/4f) mice. **G,H**) Delivery of paired-pulse electrical stimulation (2x stimuli, 50 ms interval) to thalamo-cortical afferents resulted in non-facilitating EPSCs, which did not differ between genotypes at P42 (F_(1,15)_=1.10, t_(15)_=1.047, p= 0.312, nested Student’s t-test), n=7 *Csf1r*^+/+^ (5m/2f) and n=10 *Csf1r*^ΔFIRE/ΔFIRE^ mice (6m/4f).

### Single nucleus analysis of excitatory and inhibitory neurons

To complement our electrophysiological analyses of the developing cortex we performed single nucleus RNA-seq on the P14 neocortex to determine whether any changes to the transcriptional profile of excitatory or inhibitory neurons could be detected (Fig. 6). Following an initial clustering, we selected clusters expressing the excitatory glutamatergic neuron marker Slc17a7, which encodes the vesicular glutamate transporter vGLUT, and re-clustered the excitatory neuronal nuclei (Fig. 6A). Excitatory neuronal clusters had similar abundance in *Csf1r*^ΔFIRE/ΔFIRE^ vs. *Csf1r*^+/+^ mice (Fig. 6A). Clusters were observed that expressed various cortical layer markers including Foxp2 and Tle (both layer VI), *Bcl11b*/*Ctip2* (layer V/VI), *Rorb* (layer IV) and *Cux1* (layer II/III/IV) (Fig. 6B). Moreover, differential gene expression analysis revealed no significantly changed genes (*Csf1r*^ΔFIRE/ΔFIRE^ vs *Csf1r*^+/+^), including when sex is considered (Supplementary Data 1). We also re-clustered inhibitory neurons (clusters identified by the expression of *Gad1/Gad2*). We observed that inhibitory neuronal nuclei from *Csf1r*^ΔFIRE/ΔFIRE^ and *Csf1r*^+/+^ mice clustered together (Fig. 6C), with different clusters enriched in cortical inter-neuronal sub-type markers such as *Sst, Pvalb, Vip*, *Lamp5* and *Meis2* (Fig. 6D). As with excitatory neurons we observed no significantly changed genes (*Csf1r*^ΔFIRE/ΔFIRE^ vs *Csf1r*^+/+^) including when sex is considered (Supplementary Data 2). Thus, we observe no evidence that neuronal transcriptional profiles are substantially altered in the *Csf1r*^ΔFIRE/ΔFIRE^ neocortex.

**Figure 6.**
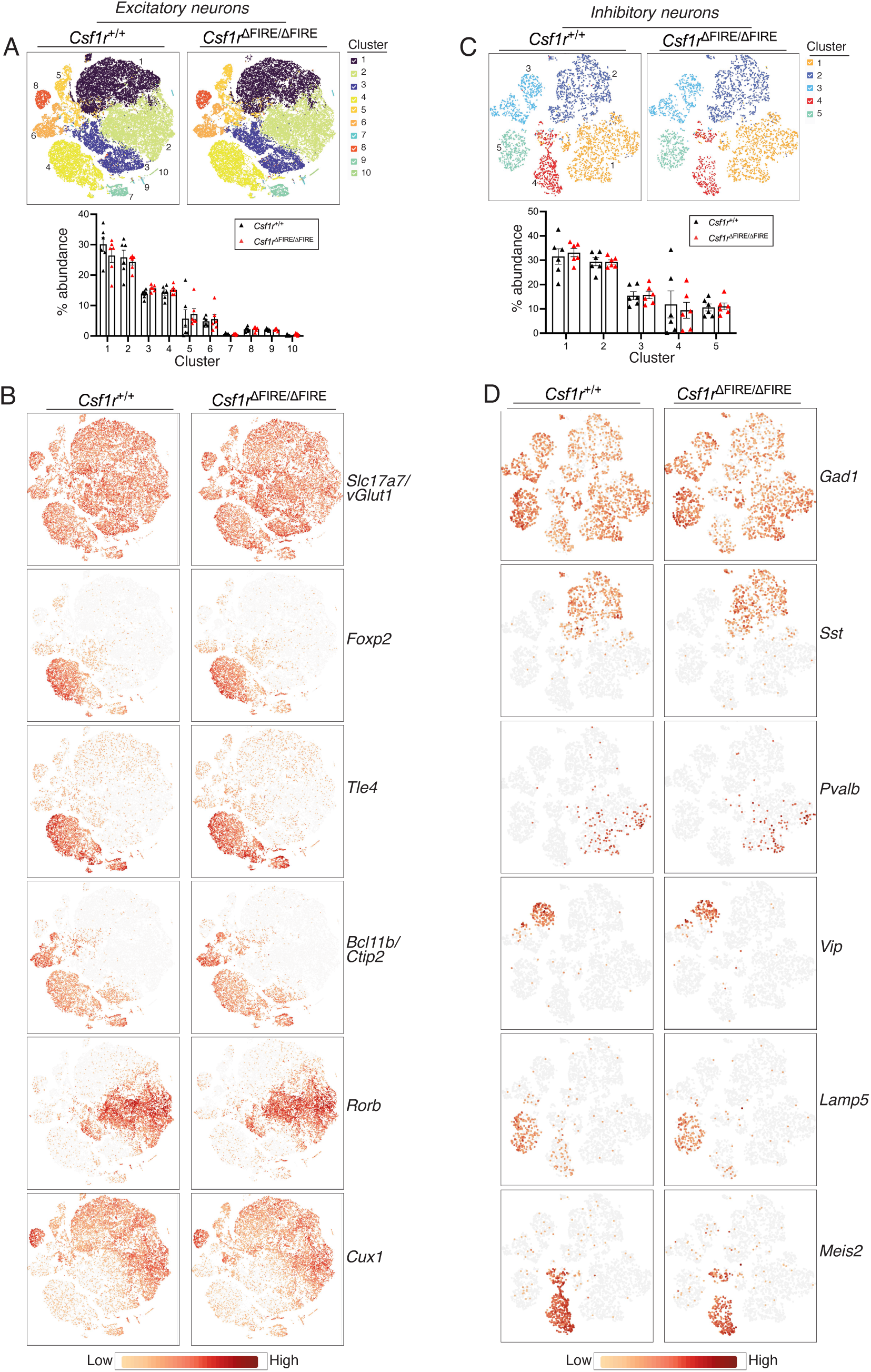
Single nucleus RNA-seq analysis of neurons from the P14 neocortex. All data are shown as mean ± SEM and all statistical tests are two-sided. **A)** Clustering of single nuclei (tSNE projection) selected based on expression of excitatory neuron marker *Slc17a7* (VGLUT1). Upper panels show clusters of nuclei whose % abundance is shown in the lower graph (n=6 per genotype). F_(1,100)_=4.3E-03, p=0.995 [genotype effect], 2-way ANOVA). **B)** The expression of cortical layer markers is compared between genotypes. Pseudo-bulk differential gene expression analysis revealed no differences in gene expression between *Csf1r*^+/+^ and *Csf1r*^ΔFIRE/ΔFIRE^ mice. **C)** Clustering of single nuclei (tSNE projection) selected based on expression of inhibitory neuron markers *Gad1* and *Gad2*. Upper panels show clusters of nuclei whose % abundance is shown in the lower graph (n=6 per genotype). F_(1,50)_=0.0004, p=0.985 [genotype effect], 2-way ANOVA. **D)** The expression of inhibitory neuron subtype markers is compared between genotypes. Pseudo-bulk differential gene expression analysis revealed no differences in gene expression between *Csf1r*^+/+^ and *Csf1r*^ΔFIRE/ΔFIRE^ mice. For A and B, n=6 *Csf1r*^+/+^ mice (3m/3f) and 6 *Csf1r*^ΔFIRE/ΔFIRE^ mice (3m/3f).

### Modest changes to astrocytes observed in Csf1r^ΔFIRE/ΔFIRE^ mice

Astrocytes may also support developmental synapse loss (in their case via MEGF10 and MERTK signaling ^31^), raising the possibility that they compensate for an absence of microglia with increased uptake of synaptic material. To investigate this, we studied the presence of presynaptic marker synaptophysin (SYP) within astrocytes in CA1 of the hippocampus at P14; which aligns with our earlier data (Fig. 1) and previous studies ^3^. We examined co-localisation of SYP and GFAP. GFAP was chosen, as basal expression is high in the hippocampus (unlike the neocortex) with strong antibody labelling, acknowledging the caveat that GFAP does not fill the entire cell. 3-dimensional confocal imaging of GFAP-positive cells revealed a similar volume occupied by GFAP-immunoreactivity in both genotypes (Fig. 7A). While we did observe the presence of SYP puncta within GFAP cells volumes (Fig. 7B, C), the number of engulfed puncta per unit volume of GFAP, including puncta that co-localised with GFAP entirely, was not different between *Csf1r*^ΔFIRE/ΔFIRE^ and. *Csf1r*^+/+^ mice (Fig. 7B). To conclude, while we do see some evidence of astrocytic uptake of synaptic material, we found no evidence of a compensatory/up-regulation of this in the absence of microglia.

**Figure 7.**
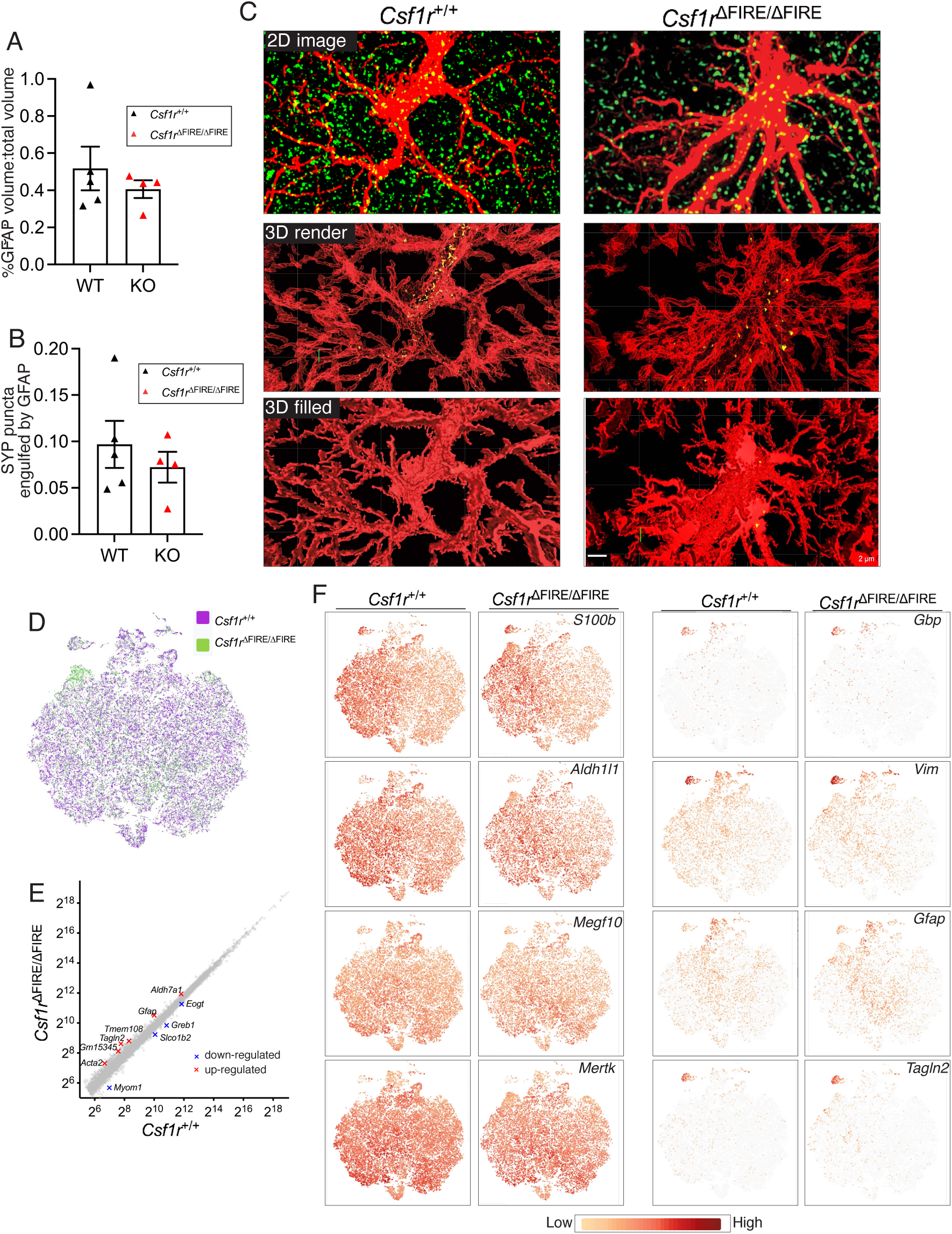
Study of astrocytes in *Csf1r*^ΔFIRE/ΔFIRE^ mice. All data are shown as mean ± SEM and all statistical tests are two-sided. **A)** Percent of total area analysed occupied by GFAP immunoreactivity in confocal stacks in CA1; t_(7)_ =0.799, p=0.451, Unpaired t-test (n=5 *Csf1r*^+/+^ mice; n=4 *Csf1r*^ΔFIRE/ΔFIRE^ mice). **B)** Synaptophysin puncta (10 nm to 1 µm in diameter) engulfed totally by GFAP immunoreactivity immunoreactivity (n=5 *Csf1r*^+/+^ mice; n=4 *Csf1r*^ΔFIRE/ΔFIRE^ mice). t_(7)_ =0.760, p=0.472, Unpaired t-test. **C)** Example images of imaged sections from *Csf1r*^+/+^ and *Csf1r*^ΔFIRE/ΔFIRE^ mice. Upper panels show raw 2D maximum projection image. Middle panels show 3D transparent render with engulfed puncta shown. Lower images show 3D surface render with engulfed puncta now not visible. Scale bar: 2 µm. **D-F)** Single cell RNA-seq of astrocytes. D) Astrocytes were sorted by FACS from the neocortex at P14 (n=4 per genotype, each 2m/2f) and subject to single cell RNA-seq (10X genomics). Small non-astrocyte contaminating cell populations were removed and astrocytes re-clustered. (D) shows the co-clustering of astrocytes from both genotypes (tSNE projection). (E) shows results of pseudo-bulk gene expression analysis: 11 genes altered out of 13,257. (F) shows expression of the indicated genes. *S100b* and *Aldh1l1* are astrocyte markers (unchanged). *Megf10* and *Mertk* are key phagocytic genes (unchanged). *Gbp* and *Vim* are reactive markers (unchanged). Examples of altered genes (*Gfap* and *Tagln2*) are shown.

To determine more generally if astrocytes are altered by an absence of microglia we performed single cell RNA-seq on FACS-sorted cortical astrocytes at P14 of *Csf1r*^ΔFIRE/ΔFIRE^ and *Csf1r*^+/+^ mice (n=4 per genotype, sex-balanced). Astrocytes from *Csf1r*^ΔFIRE/ΔFIRE^ and *Csf1r*^+/+^ mice consistently co-clustered together (Fig. 7D), and both genotypes displayed a small reactive-like cluster (enriched in reactive markers *Vim* and *Gbp2,* Fig. 8F). Differential gene expression analysis revealed only 10 genes to be altered, with only one (*Myom*) changed ≥2-fold, out of 13,257 genes meeting the expression level cut-off (Fig. 7E). A subtle increase in *Gfap* was observed as well as the reactive astrocyte-associated gene *Tagln2*, but no evidence of global astrocyte reactivity at this developmental stage (Fig. 7E, F). Examination of GFAP immuno-reactivity revealed a small increase in the cortex but not hippocampus (Extended Data Fig. 7C, D, Extended Data Fig. 7C, D). Thus, the astrocyte transcriptome exhibits quite subtle changes in *Csf1r*^ΔFIRE/ΔFIRE^ mice, although this does not rule out that they may compensate independent of changes in gene expression.

**Figure 8.**
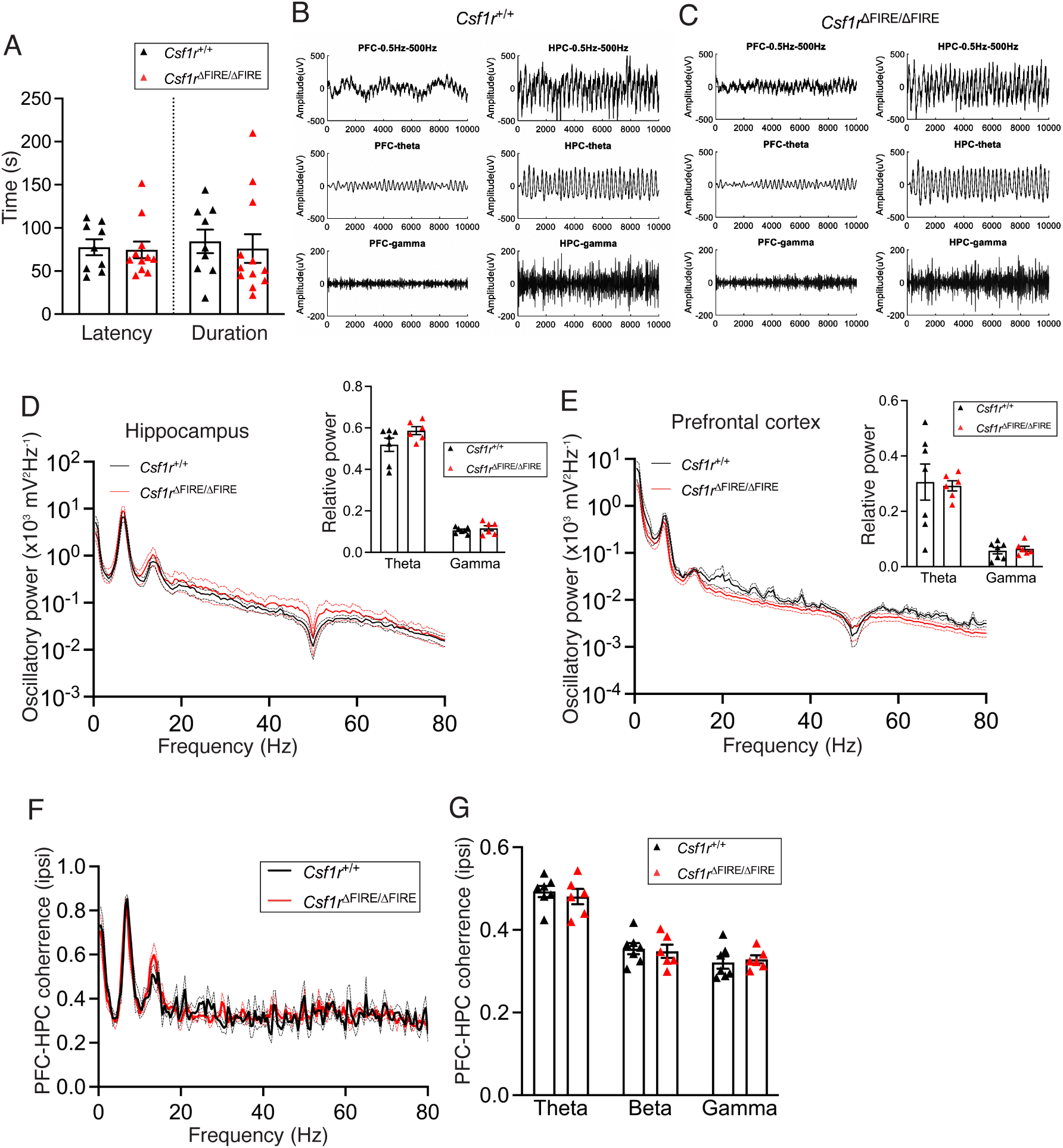
Normal seizure susceptibility and coherence of hippocampus/prefrontal cortex oscillatory activity in awake behaving animals. All data are shown as mean ± SEM and all statistical tests are two-sided. **A**) Following administration of pentylene tetrazole, we compared seizure latency (t_(18)_=0.221, p=0.828, 2-tailed Student’s t-test, p=0.882 Mann Whitney test) and total seizure duration (t_(18)_=0.233, p=0.819 2-tailed Student’s t-test, p=0.565 Mann Whitney test) between genotypes; n=9 *Csf1r*^+/+^ and 11 *Csf1r*^ΔFIRE/ΔFIRE^ mice. **B, C)** Example traces of local field potentials from the hippocampus (HPC) and prefrontal cortex (PFC) of the indicated genotypes that have been subject to a bandpass filter. **D, E**) Spectral plot of oscillatory power of LFP in CA1 of hippocampus (HPC) or prefrontal cortex (PFC) of *Csf1r*^+/+^ (black) and *Csf1r*^ΔFIRE/ΔFIRE^ (red) mice. The reduction in power around 50 Hz reflects notch filtering of the electrical line frequency. Insets: quantification of relative oscillatory power for theta (4-12 Hz) and gamma (30-80 Hz) frequency bands, normalised to total power. We observed no difference in relative power between genotypes in HPC (F_(1,22)_=3.57, p=0.072, 2-way ANOVA) or PFC (F_(1,22)_=0.009, p= 0.926, 2-way ANOVA [genotype]); n=7 *Csf1r*^+/+^ (3m/4f) and 6 *Csf1r*^ΔFIRE/ΔFIRE^ mice (3m/3f). **F, G**) *In vivo* recording of local field potentials in both CA1 of the hippocampus (HPC) and of the medial pre-frontal cortex (PFC) revealed no apparent divergence in oscillatory coherence in theta, beta, or gamma bands. F_(1,33)_ = 0.117, p=0.735, 2-way ANOVA [genotype effect]; F_(2,33)_ = 0.249, p=0.781 [genotype-frequency interaction].

### Normal seizure susceptibility and long-range activity coherence

Previous reports linking impaired microglia function and hippocampal synapse number in *Cx3cr1^−/−^* mice also reported reduced susceptibility to the pro-convulsant drug pentylene tetrazole (PTZ), attributed to a delay in microglia-dependent brain-wide circuit development ^3^. Given that development of synaptic and neuronal properties are apparently normal in *Csf1r*^ΔFIRE/ΔFIRE^ mice, we wanted to determine whether seizure susceptibility was affected. We could detect no difference in seizure latency and duration in *Csf1r*^ΔFIRE/ΔFIRE^ vs. *Csf1r*^+/+^ mice when PTZ seizures were elicited under the same conditions (Fig. 8A).

Beyond simple seizure vulnerability, neuron-microglia signalling has also been proposed to play a role in controlling long-range functional brain connectivity: young adult *Cx3cr1^−/−^* mice were reported to display a reduced coherence of local field potentials (LFP) between the medial prefrontal cortex (mPFC) and the ipsilateral hippocampus (HPC) when awake and exploring ^32^. We performed a similar analysis of HPC and mPFC LFPs in *Csf1r*^ΔFIRE/ΔFIRE^ vs. *Csf1r*^+/+^ in awake mice when exploring a virtual environment, at a similar age as previously reported ^32^. We performed spectral power analysis of LFPs recorded from the hippocampus and ipsilateral mPFC (Fig. 8D, E), and consequently assessed PFC-HPC coherence (Fig. 8F, G). We found that PFC-mHPC coherence was not different in *Csf1r*^+/+^ vs. *Csf1r*^ΔFIRE/ΔFIRE^ mice (Fig. 8F, G), suggesting that long-range connections are established normally in the absence of microglia.

## Discussion

Overall, we conclude that developing mice lacking microglia show remarkably normal synaptic density, maturation and patterning in regions where a role for these cells has been previously proposed. This conclusion is not necessarily incompatible with the previous literature reporting phenotypes of mice deficient in microglia-enriched genes such as *Cx3cr1* and *Trem2.* Rather than losing microglial function, mice deficient in *Cx3cr1* or *Trem2* may be phenotypically different due to microglia acquiring aberrant properties. For example, *Cx3cr1-*deficient microglia are morphologically different, have altered electrophysiological properties, and respond abnormally to ATP stimuli ^33^. *Trem2*-deficient microglia also have altered morphology and responses ^13,34^. It is possible that *Cx3cr1* or *Trem2-*deficient microglia have a gain-of-function effect responsible for the phenotypes observed. It is also possible that the presence of dysfunctional microglia in some way impairs the ability of the developing brain to adapt in the way that it potentially does successfully in the *Csf1r*^ΔFIRE/ΔFIRE^ mouse. Of note however, some aspects of brain development do have an absolute requirement for microglia. A recent study showed that microglia play a (largely prenatal) role in closing cortical boundaries, and that *Csf1r*^ΔFIRE/ΔFIRE^ mice exhibited a cavity at the cortico-striato-amygdalar (CSA) boundary that extended to P7 ^35^, confirming that any compensation or adaption to microglial absence does not result in a totally wild-type phenotype for developing *Csf1r*^ΔFIRE/ΔFIRE^ mice. Nevertheless, it remains a possibility that in the context of control of synapse number and patterning that astrocytes carry out functions that ordinarily are carried out by microglia. It is potentially interesting that astrocytes in *Csf1r*^ΔFIRE/ΔFIRE^ mice show a tendency to increased reactivity: we observe a small increase in Gfap expression at P14 (Fig. 7E, Extended Data Fig. 8), and by P42 our bulk RNA-seq data also shows an increase in reactive markers including *Gfap*, *Vim* and *Serpina3n*, suggesting that reactivity may increase as mice age. Whether microglia directly signal to astrocytes to prevent reactive astrogliosis is a topic for future investigation.

Another possible explanation for our findings is that the developmental functions previously attributed to microglia were based on studying mice globally deficient in *Cx3cr1*, *Cr3*, *C3* or *C1qa* ^3,4,6–8,36–38^, meaning that non-microglial processes may contribute to their phenotype. For example, Cx3cr1 and the complement system are implicated in the regulation of energy metabolism, angiogenesis, and blood-brain-barrier integrity ^15–19^, which may indirectly affect synaptic turnover. Moreover, complement is known to perform an increasing number of roles in the brain, including neural precursor proliferation and radial migration ^20–22^. Greater use of knockout models that are cell type-specific and inducible will help to attribute gene function more precisely to cell type and developmental stage.

A recent study also employed the *Csf1r*^ΔFIRE/ΔFIRE^ mouse to study aspects of neuronal development, focussing on the hippocampus ^39^. Consistent with our observations, they did not observe any change in spine density or mEPSC frequency, however, they observed a reduction in excitability, reporting that the rheobase is higher in *Csf1r^ΔFIRE/ΔFIRE^* neurons. This is a surprising finding from a biophysical point of view in the absence of a change to either AP threshold, resting membrane potential or input resistance. One difference between the studies is that the rheobase by Surala et al. ^39^ is calculated using large current steps (25 pA) relative to the rheobase (50 pA), yielding highly categorical data whereas we employed a linear ramp. The authors also reported a reduction in synaptic NMDA receptor-mediated charge transfer which we do not observe. Here the authors calculated the NMDA component of a mEPSC, rather than the usual approach of measuring AP-evoked responses (Fig 1J). Further studies are needed to rationalise these conflicting results.

Another recent study studied the role of microglia in experience-dependent maturation and function of visual circuitry ^40^. The authors used the CSF1R antagonist PLX5622 to kill microglia from P14 and found no effect on visual signaling, neuronal tuning properties in the visual cortex, and ocular dominance plasticity. The absence of a role for microglia in cortical plasticity resembles our observation in the hippocampal CA1 region (Extended Data Fig. 5) and is consistent with a previous report demonstrating that experience-dependent plasticity in the visual cortex is unchanged in Cx3cr1-deficient mice ^30^.

To summarise, the widely used term synaptic pruning implies that microglia have an active function as gardeners ^3^ in removing specific synapses to refine neurological circuits. This and other aspects of synaptic maturation are thought to require microglia to appropriately set neural circuits and connectivity. Our study demonstrates that multiple aspects of neuronal development and synapse maturation and refinement can proceed normally when microglia are absent, revealing the adaptability of brain development to absence of this cell type.

## Acknowledgements

This work was funded by the Simons Initiative for the Developing Brain and by the UK Dementia Research Institute which receives its funding from DRI Ltd, funded by the UK Medical Research Council, Alzheimer’s Society, and Alzheimer’s Research UK. We thank Jessica Valli at the Edinburgh Super-Resolution Imaging Consortium at Heriot Watt University for assistance with STED microscopy.

## Author Contributions Statement

GEH and PCK conceived the project, and directed the project along with CP, JG, AG-S. ZJ, SST, JQ, MFN, ID, DAH. M’OK, SAB, DJAW, MLi, MLiu, CH, LSdO, ML, XW, MB-L, KR, KD, CM-G, KC, TL, JQ performed the experiments and analysed the data. OD, XH and DV performed the bioinformatic analysis. GEH and PCK wrote the manuscript. All other authors provided critical feedback on the manuscript.

## Methods

### Animals

All experiments were performed under licences approved by the UK Home Office according to the Animals (Scientific Procedures) Act having been approved by the University of Edinburgh Local Ethical Review Board. This study used *Csf1r^ΔFIRE/ΔFIRE^* mice and control *Csf1r^+/+^* animals, which were generated as littermates by heterozygous *Csf1r^ΔFIRE/+^* inter-crossing. The creation and characterisation of the *Csf1r^ΔFIRE/ΔFIRE^* mice is described elsewhere ^23^. Mice are on a mixed background: 64-68% homozygous for C57BL/6J, 22-25% CBA, and 7-11% heterozygous, with approximately 2% unattributable (Mini Mouse Universal Genotyping Array). Animals were housed in communal cages (maximum 6 per cage) with ad libitum water and food in a normal 12 h light–dark cycle. Both males and females were used throughout the study. Experimenter and data analyser were blind to the genotype. Sample size was estimated utilising 3R (Reduction, Replacement and Refinement) principles, with sample size numbers calculated to achieve satisfactory power based on the variance and effect size in published data ^3,4,6–8,10,12,38^.

### Mouse pup innate motor tasks

Mice aged P3-P10 underwent two behavioural experiments to assess for motor function 1) righting reflex and 2) negative geotaxis. Each mouse completed both the righting reflex and negative geotaxis tasks every day. In between the tasks the pups were returned to heated cage to rest. When the session was over the pups were placed in their home cage and the dam was observed for signs of stress. Mice pups spent a maximum of one hour separated from the dam. *Innate righting reflex task (P3-P7):* Mice were placed on their backs on a soft platform, with their paws held together for 3 seconds. The mouse was released and the time taken for the pup to return to the prone position was recorded. A trial was defined as successful if the mouse righted themselves with all four paws on the floor. Mice were given three 15 s trials with a 20 s interval for recovery. All mice successfully managed to right themselves within the given time. *Negative geotaxis task (P3-P10):* This is the innate ability of rodents to recognise their orientation on a slope and to turn around so they are facing up hill. Mice were placed head pointing downwards on a 30% incline. Mice were placed so that all four paws were touching the surface of the incline. The pup was released and the time taken for it to turn uphill was recorded. A hand was placed approximately 10cm below the mouse to catch it if it fell. If a mouse did fall, it was replaced back on the slope in the starting position and given another trial. This was repeated until the mouse successfully reorientates itself or until a maximum of 10 attempts (9 falls) were made or until 90 s had passed. If the pup showed signs of stress such as urination or defecation, the trial ended early. The number of failed attempts and the duration to success were recorded.

### Immunohistochemistry

Mice were anaesthetised with sodium pentobarbital and transcardially perfused with ice cold PBS followed by 4% PFA in PBS. Brains were removed from the skull and post fixed in 4% PFA for 24hrs. Brains were cryo-protected in 30% sucrose overnight and then mounted on a freezing microtome in OCT compound, where coronal 50µm brain sections were prepared. Brain slices were placed in a well plate and washed with 0.1M PB and by PBS. Slices where then blocked from unspecific binding using 10% NGS, 0.3% Triton X and 0.05% Na Azide and PBS for 1 hour at room temperature. Following which, the primary antibody solution containing 10% NGS, 0.3% Triton X and 0.05% Na Azide and PBS was added (1/1000 Iba1, Abcam, ab283319) (1/500 Iba1, Wako, 019-019741) (1/1000 GFAP, Neuromics CH22102) (1/1000 Aldh1L1, Invitrogen, 702573) (1/500 Synaptophysin, Synaptic System, 102-002). Slices were incubated for 1 hour at RT followed by 24h at 4°C. Well plate was removed and allowed to come to temperature before the secondary antibody solution was added (10% NGS, 0.3% Triton X and 0.05% Na Azide and PBS) for 24 hrs. Following which slices were washed with PBS and 0.1PB for 1 hour before mounted on glass slides and cover slipped. Images were obtained using the Leica SP8 confocal microscope. 1024×1024 pixel images of the CA1 (Bregma A-P: –1.75 to –1.9 mm) and the dLGN (Bregma A-P: –2.25 to –2.5 mm) were taken using the 10x (N.A. 0.45) and 20x (N.A. 0.8) objectives. Multiple images were taken for each animal. To analyse astrocyte density images were opened using ImageJ software. For cell counts, a counting grid composed of 200 x 200 µm squares was superimposed over the stratum radiatum of the CA1. Using the cell counter tool the number of DAPI, GFAP and ALDH1L1 positive cells were quantified. Each DAPI, GFAP and ALDH1L1 cell was counted only if the cell body was within the grid and over 50% of the protrusions were within the grid. The average cell density was measured for each animal and reported as the number of GFAP and ALDH1L1 per 100 µm^2^ and as percentage of total DAPI cells.

### Bulk and single nucleus RNA-seq

Bulk tissue RNA extraction was carried out as described ^23^. Briefly, brains from saline perfused mice were dissected, snap frozen, and subsequently disrupted in the Precellys24 Homogenizer® (Bertin Instruments). RNA isolation was performed using the RNeasy Plus Mini kit (QIAGEN). RNA-sequencing reads were mapped to the primary assembly of the mouse reference genome using the STAR RNA-seq aligner 2.7.11 ^41^. Tables of per-gene read counts were generated from the mapped reads using featureCounts ^42^. Differential gene expression was then performed in R using DESeq2 ^43^. For bulk RNA-seq 6-7 mice per genotype was analysed. To sort astrocytes (4-5 mice per genotype), mouse neocortices were harvested then dissociated with Adult Brain Dissociation Kit (Miltenyi Biotec) on a gentleMACS Octo Dissociator using program 37C_ABDK_01. Dissociated samples were then treated with debris and red blood cell removal steps to obtain cell suspensions. For FACS, cells were resuspended in a final volume of 100 uL 0.1% PB buffer and incubated with fluorescent conjugated antibodies at 4° C for 30 minutes. The following antibodies were used: ACSA2 APC (1:200, Miltenyi Biotec, 130-116-245) and O4 PE (1:100, Miltenyi Biotec, 30-117-357). ACSA2 APC (+) and O4-PE (−) live single cells were selected as the astrocyte population. These were then loaded onto a 10x Chromium Controller. For single nucleus-RNAseq, neocortices were acutely collected, flash-frozen, and then stored at –80°C. Nuclei isolation kit (Sigma-Aldrich; NUC201-1KT), DTT (R0861), RNAse inhibitor (Life Technologies; AM2694), 30 µm cell strainer (Partec CellTrics), and 30 G insulin needles (BD; 324826) were used. Cryopreserved tissue samples were thawed on ice. Lysis was performed with PURE buffer, 0.1M DTT, 10% Triton, and RNAse inhibitor. The samples were mechanically dissociated in lysis buffer using a P-1000 pipette 10 times, and further 3 times with insulin needles. The lysate was mixed with sucrose cushion, which was prepared using PURE 2M sucrose, sucrose cushion buffer, 0.1M DTT, and RNAse inhibitor. This mixture was then filtered through a 30 µm cell strainer. In a new low-binding tube, 200 µl of sucrose cushion was added and carefully overlaid with 560 µl of the filtrate. The samples were centrifuged for 45 minutes at 4°C. The supernatant was discarded, and the pellet was resuspended in Dulbecco’s phosphate-buffered saline (DPBS) (14190-094) + 0.5% BSA with RNAse inhibitor. The suspension was then centrifuged at 1000 G, repeated, and finally resuspended in 200 µl DPBS + 0.5% BSA with RNAse. Sorting of nuclei (6 mice per genotype) was done by flow cytometry. The gating was set based on size and granularity using Forward Scatter (FSC) and Side Scatter (SSC) to capture singlets and remove debris. Nuclei were stained with DAPI for detection. Quality control was conducted using Luna FX7 with Acridine Orange/Propidium Iodide (AO/PI), and the input nuclei number was 20,000 per sample. For library preparation, nuclei suspensions were loaded onto a 10x Chromium Controller (10x Genomics). Single-nucleus transcriptomic amplification and library preparation were conducted with the Chromium Single Cell 3ʹ v3.1 Reagent Kit (10x Genomics). Libraries were first sequenced on the iSeq 100 System (Illumina Inc, #20021532) using the iSeq 100 i1 Reagent v2 (300 cycle) Kit (#20031371). Library molarity for sequencing was calculated using the Qubit dsDNA quantification results and the fragment size information from the Bioanalyser results. Libraries were normalised to 10 nM and equal volumes were pooled and diluted for sequencing. PhiX Control v3 (#FC-110-3001) library was spiked into the run at a concentration of ∼4% to help with cluster resolution and facilitate troubleshooting. iSeq data was then analysed to so that pools could be rebalanced for subsequent deep sequencing. Deep sequencing was performed on the NextSeq 2000 platform (Illumina Inc, #SY-415-1002) using the NextSeq 1000/2000 P3 Reagents (100 cycles) v3 Kit (#20040559). PhiX Control v3 (#FC-110-3001) library was spiked in at a concentration of ∼1%. For single-cell and and single-nucleus sequencing data, sequencing reads were mapped to the mouse genome, and per-cell, per-gene count matrices produced, using 10x CellRanger version 7.0.1 ^44^. QC, normalisation and clustering of data was performed using the Seurat R package, version 4.4.0 ^45^. For single-nucleus sequencing, ambient RNA was estimated and removed using the SoupX R package version 1.6.2 ^46^. Doublets were identified and removed using the ScDblFinder R package, version 1.10.1 ^47^. Pseudo-bulk differential expression analysis was performed by summarising single cell gene expression profiles at the subject level using the aggregateBioVar R package, version 1.6.0 ^48^, then differentially expressed genes between genotypes were calculated using DESeq2 version 1.36.0.

### Slice preparation, patch-clamp and field potential recordings

Acute slices were prepared as previously described ^49^. Once cut, slices were transferred to a holding chamber containing carbogenated sucrose-aCSF (whole-cell recordings) or aCSF (field recordings). Slices were allowed to recover for ≥ 30 minutes at 35 °C until needed. For electrophysiological recordings, slices were transferred to a submerged recording chamber, perfused with carbogenated aCSF ^49^ at a flowrate of 3-6 ml/min at 30-31 °C.

For whole-cell patch-clamp recordings, slices were placed in the recording chamber of an upright microscope (SliceScope, Scientifica, UK) and visualised using infrared differential interference contrast (IR-DIC) microscopy. For CA1 recordings, a stimulating electrode placed in *stratum (str.) radiatum.* Cells were chosen under high magnification within *str. pyramidale* as having large ovoid somata, with a clear apical dendrite entering *str. radiatum*. For layer 4 (L4) of the primary somatosensory (S1) cortex, stimulating electrodes were placed in either the ventro-basal thalamus (VBT) or internal capsule. L4 was identified based on the presence of barrel-like structures observed under low-magnification. For whole-cell recordings, borosilicate glass microelectrodes were pulled on a horizontal electrode puller (model P-87 or P1000, Sutter Instruments), which were filled with either Cs-gluconate based (in mM: 140 Cs-gluconate, 4 CsCl, 0.2 EGTA, 10 HEPES, 2MgATP, 2 Na_2-_ATP, 0.3 Na_2-_GTP, 10 Na_2-_ phosphocreatine, 2.7 biocytin, and 5 QX-314; pH 7.4, 290–310 mOsm), or a K-gluconate (in mM: 142 K-gluconate, 4 KCl, 0.5 EGTA, 10 HEPES, 2 MgCl_2_, 2 Na_2_ATP, 0.3 Na_2_GTP,10 Na_2_phosphocreatine, and 2.7 biocytin; pH 7.4, 290–310 mOsm) based internal solutions, which gave a tip-resistance of 5-6 MΩ. Signals were recorded using a Multiclamp 700B amplifier, digitized using a Digidata 1550b (10 or 20 kHz). Cells were rejected if they required a holding current of >200 pA to maintain voltage clamp at –70 mV, if series resistance started at >30 MΩ, or if series resistance changed by >20% over the course of the recording. Miniature and evoked synaptic recordings were carried out following wash-in of the Cs-gluconate internal solution for 2-5 minutes at a holding potential of –70 mV. Miniature EPSCs were recorded in the presence of 300 nM tetrodotoxin (TTX). For measurements of NMDA/AMPA ratio, CA1 pyramidal neurons or S1 L4 neurons were recorded in the presence of 50 µM picrotoxin. Evoked PSCs were generated using a twisted Ni:Chrome bipolar wire connected to either a constant-voltage or constant-current stimulator (Digitimer, UK). In hippocampal slices, the stimulus intensity was adjusted as to produce a monosynaptic AMPA receptor-mediated EPSCs of approximately 200 pA. For S1 recordings, an extracellular field electrode (a patch pipette filled with ACSF) was first placed in a L4 barrel, and electrical stimuli delivered to identify synaptic connections. The stimulating electrode was moved between consecutive barrels until a synaptic response was identified. After identifying connectivity, neurons were then recorded in that barrel. Monosynaptic AMPA receptor-mediated EPSCs were generated (WT_median_= –86 pA [Range: –17 to-491 pA]; *Csf1*^+/FIRE^_median_= –68 pA [range: –8 to –346 pA]). EPSC amplitudes were measured as the average peak amplitude (over a 2ms average) of 10 responses, elicited 20 s apart. Assessment of NMDA receptor function was performed at + 40 mV. For mixed AMPA/NMDA EPSCs the mean response 50-60 ms post-stimulus onset, where the AMPA receptor response has completed its decay, was taken as a proxy of NMDA response amplitude. Pharmacologically isolated NMDA responses were obtained in the presence of by washing on 50 µM CNQX until the AMPA response was completely abolished. For GABA_A_ receptor-mediated IPSCs, neurons were held at 0 mV in voltage-clamp.

The intrinsic properties of neurons were measured using a K-gluconate internal solution. Resting membrane potential (RMP) of the cell was recorded with current clamped at 0 pA, and all other protocols recorded with appropriate current injection to hold the cell at −70 mV. Input resistance and membrane time-constant were calculated by injecting a −10 pA step. Current-frequency responses were assessed using a series of 500 ms rectangular current injections ranging from 25-400 pA (25 pA steps). Rheobase was assessed using a linear ramp from –100 – 400 pA over the course of 2s. Analysis of whole-cell recordings was performed using the open source software package Stimfit, Clampfit, or custom-written MATLAB scripts (see Supplementary Software 1 and 2). Following recording, neurons were re-sealed by generating out-side out patches, then fixed with 4% paraformaldehyde for subsequent visualisation and immunohistochemistry.

For hippocampal field EPSP (fEPSP) recording, the CA3 region was removed. FEPSPs were elicited by delivering a short pulse of electrical current (0.1 ms) to Schaffer Collateral axons through twisted Ni:Chrome bipolar wire connected to either a constant voltage or constant current stimulator. Borosilicate glass recording microelectrodes with a resistance ranging from 1–3 MΩ were pulled and filled with aCSF. Stimulus intensity was set to 50% of the maximal fEPSP amplitude. LTP was induced by high-frequency (two trains of 1s 100 Hz stimulation, 20s inter-train interval) following ≥20 minutes of stable baseline. Data was amplified through a EXT-02B amplifier (npi electronic GmbH) and digitised at a rate of 20 kHz through a BNC-2090A (National Instruments). Data acquisition and analysis were performed on WinLTP ^50^.

### STED microscopy

Hippocampal slices from P28 mice were prepared for slice physiology and individual CA1 pyramidal cells were filled with biocytin using a patch. Slices were incubated at 4^0^C overnight in solution containing streptavidin conjugated to Abberior STAR 580 (1:500) (Abberior STAR 580, ST580), 3% NGS, 0.1% Triton=X, and 0.05% Na-Azide. Slices were washed with 0.1M PB, mounted onto glass slides. Time gated STED (gSTED) images were obtained using a STED microscope (Leica SP8 3X gSTED). Individual apical oblique dendrites from CA1 pyramidal cells were imaged using a high magnification objective (93x glycerol immersion objective lens, N.A 1.3). Sections were illuminated with 580nm light. Image stacks of apical oblique dendritic sections within the stratum radiatum of the CA1 were imaged using a step size of 0.09µm. The image stacks were Nyquist sampled with a pixel size of 20nm and at a scan speed of 200 Hz using six-line averaging. STED images were deconvolved using Huygens’s STED module (Huygen’s STED, Scientific Volume Imaging, Netherlands). Measurement of spine morphological parameters were carried out using ImageJ. For WT animals 124 spines from 5 animals were measured. For KO animals, 138 spines from 5 animals were measured. 2 dendritic sections were imaged per cell and 1-4 cells were imaged per animal. Statistics were performed on the number of independent biological replicates (i.e. animals).

### Labelling, imaging and analysing contralateral and ipsilateral retinal inputs into the dLGN

P4-5 and P10-11 were anesthetized with isoflurane and decapitated. The skin was removed from the head and the snout clipped. The cornea was peeled back to expose the inner part of the eye. Humorous jelly was removed exposing the retina and optic nerve. Crystals of lipophilic dye (DiD, Thermoscientific, D7757, or DiI, Biotium, 60010) were placed inside the eye to differentially label the contralateral and ipsilateral eye and the cornea was replaced. The head was stored in 4% PFA and incubated at 37^0^C for a week. PFA was then removed and replaced with PBS and 0.1% Na Azide and incubated at 37^0^C for a further eight weeks.

Following incubation, the head was removed and placed in 4% PFA overnight at 4^0^C. The brain was removed from the skull and 50µm slices were prepared and mounted. Images of the dLGN were obtained using Zeiss LS800 confocal microscope using the 20x objective (N.A. 0.8). 1024×1024 pixel and tiled together. All images were acquired and analysed blind to genotype (Bregma A-P: –2.25 to – 2.5 mm). Image analysis was performed using the FIJI package of ImageJ ^51^. The ventral region of the LGN (vLGN) receives no contralateral input from the retina, and as such the vLGN should have a fluorescence level similar to that of background. If images failed to meet this criterion they were omitted from analysis. Images were opened in ImageJ software and split into two channels (ipsilateral and contralateral). Background fluorescence was removed using the ‘rolling ball radius filter; set to diameter of 200 pixels. The outline of the dLGN was drawn using the freehand tool, this excluded the vLGN and the optic tract. This outline was saved using the ‘region of interest’ (ROI) manager tool. The threshold of the contralateral region was set for multi-threshold levels, 0-150 (in 5-pixel steps). A mask was created for each threshold step and saved to the ROI manager. A threshold of 35-45 was set for the ipsilateral image, ensuring the entire ipsilateral region was included and a mask was created using the selection tool and saved to the ROI. Using the ‘and’ option of the ROI tool, the mask of the ipsilateral region was overlaid with the contralateral mask and the level of overlap was measured in pixels for each threshold. The fraction of ipsilateral region without overlap was then calculated for each contralateral threshold and reported as the fraction of segregated ipsilateral inputs. Segregation curves were plotted for each genotype and age.

### Barrel cortex anatomical analysis

P10 mice were sacrificed and transcardially perfused with ice cold PBS followed by 4% PFA in PBS. Brains were removed from the skull and post-fixed in 4% PFA for 24h. 50µm brain sections were prepared on the tangential plane. Two different flattening techniques were used to analyse a) barrel area of PMBSF and b) relative position of PMBSF. a) The two hemispheres were separated, and the thalamus, hippocampus, entorhinal cortex, and striatum were removed.. Tissue is placed on a freezing block of OCT on the freezing microtome and gently flattened with a glass coverslip and 50µm sections were prepared. b) The two hemispheres were separated, and the thalamus, hippocampus and part of striatum were removed. Tissue was placed between glass slides with capillary tubes as spacers and flattened. The tissue was fixed in 4% PFA for 24 hours before processing. The flattened hemisphere is placed on the freezing microtome and cut (Bregma: 1.75 to –1.9 mm). To perform immunohistochemistry, brain slices were placed in a well plate and washed with 0.1M PB and by PBS. Slices where then blocked using 10% NGS, 0.3% Triton X and 0.05% Na Azide and PBS for 1 hour (RT). Following which, the primary antibody (1/1000 VGLUT2, 2B scientific, 135-402-SY) solution containing 10% NGS, 0.3% Triton X and 0.05% Na Azide and PBS was added. Slices were incubated for 1 hour at RT followed by 24hrs at 4 degrees. The well plate was removed, primary antibody solution removed (plus 3X PBS washing) and allowed to come to temperature before the secondary antibody solution (10% NGS, 0.3% Triton X and 0.05% Na Azide and PBS) was added for 24 hrs. Following this, slices were washed with PBS and 0.1 PB for 1 hour before being mounted.

### Analysis of dendritic spines and neuronal morphology

CA1 pyramidal cells were filled with biocytin during electrophysiology experiments via the patch pipette. Slices were removed from the recording chamber and transferred to a 24-well plate containing 4% PFA for approx. 1 hour after which they were transferred to PBS. Cells filled with biocytin were stained for streptavidin. Secondary antibody solution made of 3% NGS, 0.1% Triton-X, 0.05% Na-Azide and Streptavidin Alexafluor 488 (1:500), was added and incubated for 24 hours at 4^0^C. Slices containing streptavidin labelled CA1 pyramidal cells were imaged using the Zeiss 1800 confocal microscope. Images were acquired and analysed blind to genotype. For neuronal morphology analysis individual pyramidal cells were imaged using 20x objective (N.A. 0.8) Z stacks of 2µm step size were imaged along the length of the neuron and tiled together using imageJ. For spine density analysis, sections from three different dendritic compartments (apical oblique, apical tuft and basal) were imaged. Two to three dendrites were imaged for each compartment. Images were acquired with a 63x oil emersion objective (N.A 1.4) using Nyquist sampling. For a section of dendrite two to three Z stacks with a step of 0.12µm were taken along the dendrite.

For neuronal morphology analysis a 3D neuronal reconstruction of each cell was created by stitching images together using the tiling function of ImageJ. Images were imported into Huygens Essentials software package (Scientific Volume Imaging, The Netherlands, http://svi.nl), and deconvolved using the fast CMLE algorithm, with SNR:20 and 100 iterations. Neuronal reconstructions were created of CA1 pyramidal cells. Tiled z-stacks were imported into ImageJ and the Simple Neurite Tracer (SNT) plug-in ^52^ was used to create SWC trace files for each cell. Using the tracing tool, the length of dendrites was traced from the cell soma. By scrolling through the z-stack, individual dendrites can be traced as they transverse through the slice. Trace files were converted into a 3D neuronal reconstruction representing individual cells. Neuronal reconstructions can be used to analyse dendritic complexity.

Neuronal morphology was assessed using Sholl analysis ^53^. The number of dendritic intersections is quantified every 20 µm concentric distance from the cell soma. Using the SWC trace files, new trace files were created to account for the total number of dendrites and for each dendritic compartment. A Sholl analysis was performed using each of these trace files and the SNT. The total number of dendritic branching at a given distance from the soma was reported. To quantify the total length of dendrites within a given compartment as path order analysis was carried out. This term, ‘path order’, refers to the hierarchy of dendrites originating from the soma in respect to their branch order, primary, secondary, tertiary. The ‘path order’ tool of SNT was used to produce a table comprising of the numbers of primary, secondary, and tertiary dendrites, and the length of each dendrite. Any dendrites above the tertiary hierarchy were recorded as tertiary dendritic paths. The total length and number of dendrites within each hierarchal compartment was reported.

For spine density analysis, deconvolved images of dendritic sections were opened in ImageJ and the density of dendritic spines from apical oblique, apical tuft and secondary basal dendrites was quantified using “Cell Counter” plug-in on ImageJ. The number of protrusions (spines) were manually counted from either side of the dendritic shaft starting from 5-10 µm from the branch point. The length of the dendritic section was measured and recorded using the ‘Measure’ tool.

### Calculating astrocyte uptake of synaptic material

Images of the CA1 stratum radium were obtained using the Leica SP8 confocal microscope. Image stacks were acquired using Nyquist sampling with a z step size of 0.12 µm and pixel size of 45 nm. Z-stack images of CA1 stained with antibodies against Synaptophysin (SYP) and GFAP were first pre-processed: digital images were first normalized to standardize intensity scales across the dataset. The synaptophysin channels were clarified using a Particle Sharpen filter from OpenCV (Python package), while GFAP channels were smoothed with a Discrete Gaussian Blur, reducing random noise and enhancing meaningful signal. Image classification was conducted using a Random Forest algorithm in Vision4D, tailored through training based on 15 manually annotated sections each of Synaptophysin, GFAP, and background signals. This model focuses on pixel-wise classification, relying on local contextual information without considering the overarching morphology of the labeled structures.

Segmentation outputs defining GFAP and Synaptophysin were encoded into TIFF format for in-depth analysis using Python version 3.9. To address z-stretching artifacts, operations from ‘scipy.ndimag’ were employed, including ‘binary_erosion’ with ‘generate_binary_structur’ to discern connected objects in the z-plane based on a parameterized structuring element. Minor GFAP elements below a volume threshold of 10 µm³ were excluded as noise using ‘binary_closing’. Synapses were filtered based on their cubic volume to between 0.01um^3 and 1um^3, based on previous quantitation of presynaptic bouton size in CA1 ^54^.

For quantification of engulfment, 3D mesh models of GFAP and Synaptophysin structures were generated using ‘marching_cubes’ from ‘skimage.measur’, and surface areas were determined by ‘mesh_surface_areà. The ‘regionprops’ function provided dimensional and positional descriptions of labeled regions, facilitating the analysis of synapse size and spatial relationships. Overlapping volumes between Synaptophysin and GFAP were quantified using ‘distance_transform_edt’ from ‘scipy.ndimag’, identifying engulfment. Synaptic engulfment (100% engulfed) were then associated and analyzed in terms of relative GFAP volumes using spatial overlap metrics. Results were stored and manipulated with ‘pandas’ DataFrame structures, which facilitated data handling and subsequent statistical testing.

### Induction of seizures

P17-P20 mice were in injected intraperitoneally (i.p) with the proconvulsant drug Pentylenetetrazol (PTZ) (MERK, P6500) at a dose of 70 mg/kg, 10ml/kg. Mice were immediately placed into a 60cm x 60cm arena and recorded for a maximum of 10 mins or when they reached a maximum severity level as assed by the modified Racine scale (Lüttjohann et al. 2009). Each severity level is characterised by a behaviour and associated with a number, 0-7. A maximum time of 10 mins was given and if the animals did not reach level 7 at this stage they were removed and culled. No mice reached the maximum time. Experimenter was blind to genotype during injection and analysis. For analysis, videos were opened and analysed using BORIS software. The latency to first seizure and inter seizure interval time was recorded. The duration of each seizure was measured and the severity level was recorded.

### In vivo electrophysiology

3-5 months old *Csf1r*^ΔFIRE/ΔFIRE^ mice and their littermate controls were anesthetized with isoflurane (3 % induction, 1% maintenance, 0.5L/min in pure oxygen) and mounted into a stereotaxic frame. 2 steel screws (1 mm diameter) were placed in small boreholes drilled above the cerebellum and olfactory bulb. A custom-made head-ring for later head-restraint training and recording was attached to the skull and the screws and tightly fixed using bone cement (Refobacin, Biomet). After the surgery, mice were injected with Carprofen (Rimadyl, 120-130 µL S.C., 20 mg/kg) and received a liquid supplement with 0.9% NaCl, 200-300 µL I.P. Mice were given at least 7 days to recover. Mice were then handled and habituated to head-restrained on an air-cushioned styrofoam ball in a virtual reality (VR) system (JetBall, Phenosys). After habituation, mice were water-restricted (approximately 1.2ml water/day) and trained to run in a VR linear treadmill for liquid rewards. When the animals could perform sufficient runs on the treadmill after 3-4 days, they were anesthetized, and three small craniotomies were drilled. One above the prefrontal cortex (AP:≈ 1.9 mm; ML:≈ 0.5-1 mm) and two above the hippocampus in each hemisphere (AP: ≈ –1.9 mm; ML:≈ ± 1.5 mm). Dura was removed. Craniotomies were sealed and protected by silicone (Kwik-Cast, World Precision Instruments). Animals were left for recovery for at least 8 hours before electrophysiological recordings started.

To perform the virtual reality linear track treadmill task, mice were head-fixed on an air-cushioned styrofoam ball within a 270° surround TFT monitor system (JetBall, PhenoSys). Mice were trained to run to advance in a custom-made virtual linear corridor. The running path length was 50 cm for each trial. Once the animal reached the end of the virtual corridor in each trial, a 50 µL liquid reward (5% sucrose water) was delivered through a spout. After a 3-second inter-trial interval, animals were teleported to the start of the virtual corridor to initiate the subsequent trial. Each animal was trained once daily for 15-30 minutes for at least 3 days when sufficient amounts of liquid (≈ 1-1.2 mL) were obtained during the task. Trial signals in the tasks were recorded as TTL pulses by the acquisition device. The animal’s locomotion was detected by an XY-motion sensor at 50Hz.

To perform recordings, mice were attached to the stereotaxic frame in the VR and were allowed to perform the linear treadmill task. Two silicone probes, each consisting of 128 channels on 4 parallel shanks (A4X32-Poly2-5mm-23s-200-177, NeuroNexus), were inserted into the prefrontal cortex and the hippocampus, respectively through the craniotomies prepared. Local field potential (LFP) signals were recorded using an Intan 128-channel head-stage from each probe and an Intan RHD recording controller. Signals were sampled at 20 KHz and digitized as 16-bit signed integers. All signal processing and data analysis were performed in Matlab (Mathworks). The power-line interference (50, 100, 150, and 200Hz) was removed from the LFP signals by a 2^nd^ order band-stop filter. After eliminating further noises, LFP signals were down-sampled to 2 kHz, using a resample function which applies an FIR antialiasing low-pass filter. Then LFPs were filtered between 0.5 and 500 Hz using a 2^nd^-order Butterworth band-pass filter. Segmentations of LFP data where the animal’s running speed was 5–10 cm/s were extracted for analysis. A 5-s window, backwards from 0.5 s prior to the reward delivery time, was selected in each trial. The power spectrum density and coherence analysis were obtained by the pwelch function (window: 2s, overlap: 50%) and the mscohere function (window: 2s, overlap: 50%), respectively. A theta (6-10Hz)-to-delta (2-4Hz) ratio > 4 of mean power spectral density in the hippocampus was applied to select clear running/alertness-associated theta epochs as described previously ^55–58^. Power was calculated as the sum of the power spectral density within each frequency band, and coherence was calculated as the mean within each frequency band (Theta: 4-12Hz; Beta: 15-25Hz; Gamma: 26-70Hz).

### Statistics

Throughout the manuscript the statistical test is stated, along with p values, and test statistic. Statistical tests performed are always two-sided, and performed on biological replicates, not technical replicates. Multiple slices or cells measured from the same animal were treated as technical replicates. Data were generally assumed to be normal, but when failing normality test a non-parametric version of the t-test (Mann Whitney) was used. T-tests included Welch’s correction but otherwise data variance was assumed to be similar. Differential expression analysis of RNA-seq data was performed using DESeq2 88, with a significance threshold calculated at a Benjamini–Hochberg-adjusted P value of <0.05. Sample sizes ranged from 4 to 15 animals per condition/genotype. For cell counting and recording the experimenter was blind to the condition or genotype. For all images displayed, contrast and brightness are applied in a linear manner across the entire image.

## Code availability

Custom MATLAB scripts Supplementary Software 1 and Supplementary Software 2 are supplied as supplemental material.

## Data availability

Accession code relating to RNA-seq data (ArrayExpress) are E-MTAB-14156 and E-MTAB-14160. Microglia-enriched genes employed data as described ^59^ and publicly available (SRA accession SRP135960). Data that support the findings of this study are either either available in the paper and its supplements, or available from the authors on request.

---

**Extended Data Fig. 1.**
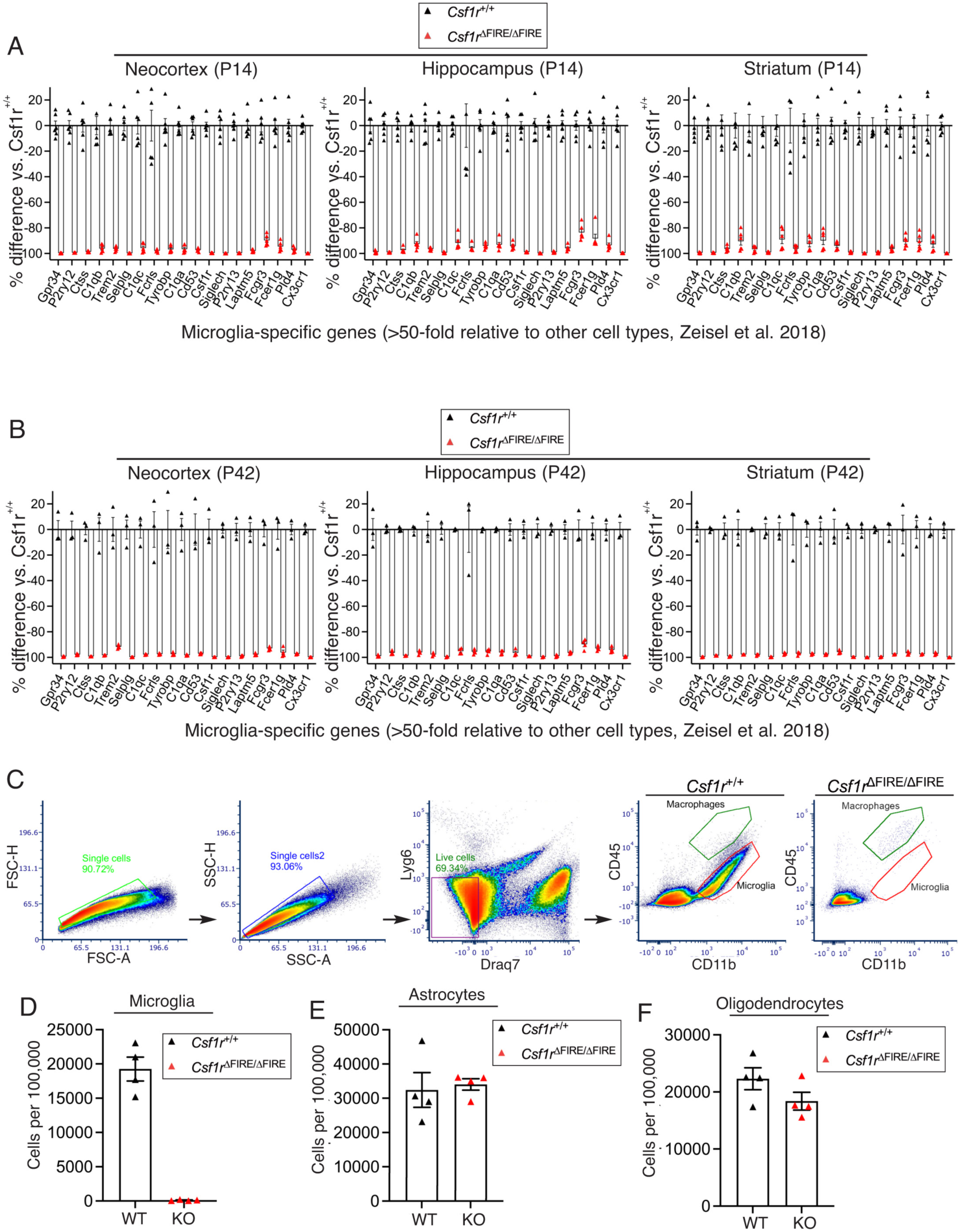
Molecular and anatomical characterisation of Csf1r^ΔFIRE/ΔFIRE^ mice during early development. All data are shown as mean ± SEM and all statistical tests are two-sided **A,B)** RNA sequencing data showing the relative expression of microglia-specific genes in the Csf1r^ΔFIRE/ΔFIRE^ compared to Csf1r^+/+^ for neocortex (left), hippocampus (middle) and striatum (right), at P14 (A) or P42 (B). Genes were selected based on previously published single-cell sequencing data (Ziesel et al., 2018) that described 265 subclusters in the CNS and PNS. To define microglia-enriched genes, we calculated the fold-enrichment of expression in the “MGL1” subcluster relative to the next highest expressing non-immune cell subcluster, requiring an average of ≥1 FPKM in wild-type mice. Data relating to genes with a microglial enrichment of ≥50-fold are shown. Sample size at P14: 6 Csf1r^+/+^ mice, 6 Csf1r^ΔFIRE/ΔFIRE^ mice (3 males and 3 females for both). Sample size at P42: 3 Csf1r^+/+^ mice, 4 Csf1r^ΔFIRE/ΔFIRE^ mice. **C-F)** Cells from the neocortex of P14 mice of the indicated genotypes were analysed by flow cytometry (see Methods). **(C)** shows the sequential gating strategy and example plots of CD11b (x-axis) and CD45 (y-axis) illustrating the loss of CD11b^+^/CD45^low^ microglia in mice, with far less abundant CD11b^+^/CD45^high^ border-associated macrophages retained, (D-F) shows quantitation of microglia, astrocytes and oligodendrocytes (n=4 per genotype).

**Extended Data Fig. 2.**
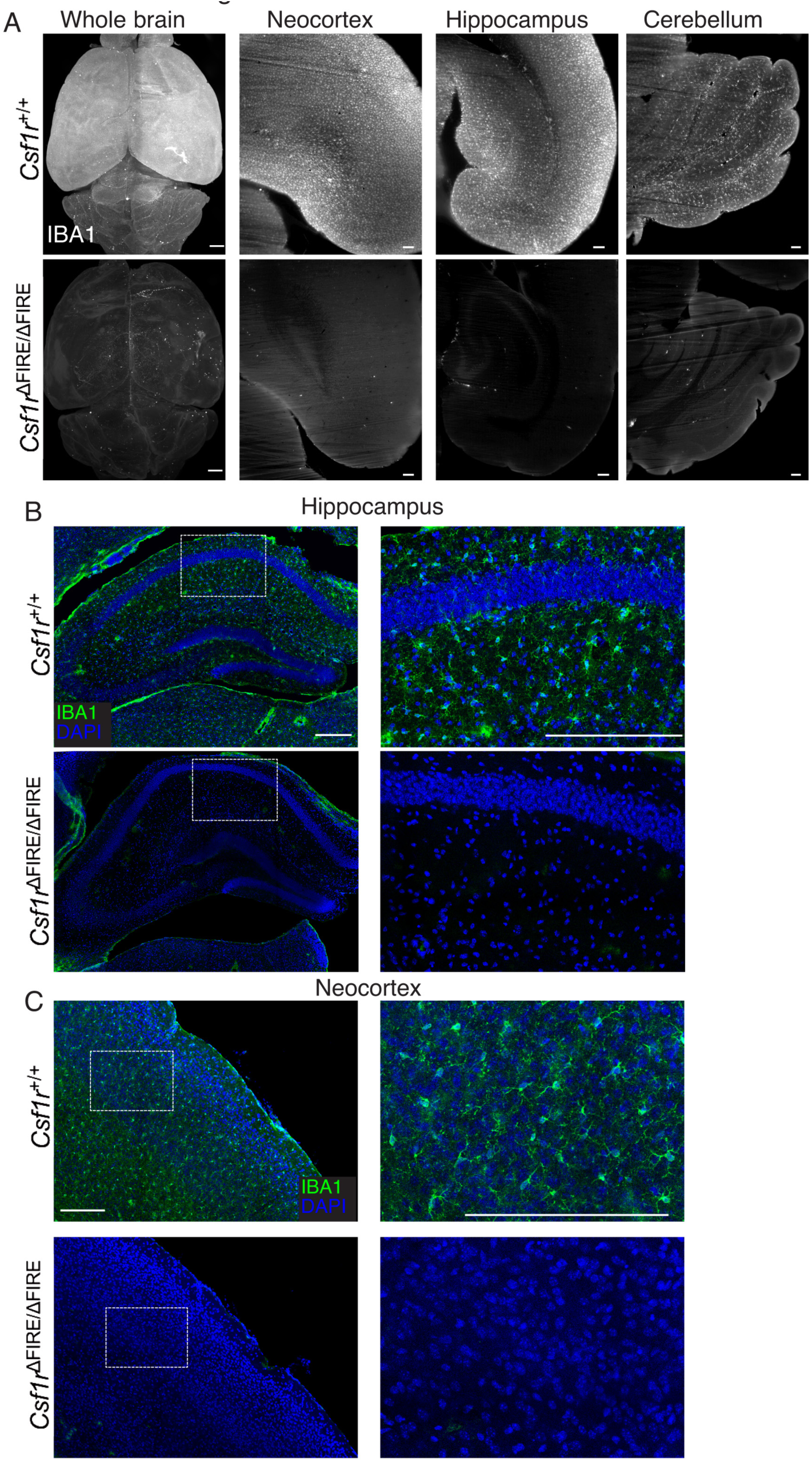
Loss of IBA1-positive microglia in the *Csf1r*^ΔFIRE/ΔFIRE^ mouse. **A**) Example images (representative of and 7 *Csf1r*^+/+^ and 6 *Csf1r*^ΔFIRE/ΔFIRE^) from whole-brain imaging performed using iDISCO. Brains were labelled for IBA1 (greyscale) for *Csf1r*^+/+^(upper) and *Csf1r*^ΔFIRE/ΔFIRE^ (lower) mice at P28. Flattened volume images are shown for the whole brain (left), neocortex (middle left), hippocampus (middle right), and cerebellum (right). Note the prominent labelling for Iba1 in *Csf1r*^+/+^ brains, but not *Csf1r*^ΔFIRE/ΔFIRE^ brains. Scale bars: 500 µm (left, whole brain), 100 µm (middle, right, sub-regions). Note that the tiny amount of residual IBA1 staining in the *Csf1r*^ΔFIRE/ΔFIRE^ brains is due to border-associated macrophages ^1,2^. **B,C)** Conventional immunohistochemical analysis of IBA1 expression (green), counterstained with DAPI, in the indicated brain regions of *Csf1r*^+/^ and *Csf1r*^ΔFIRE/ΔFIRE^ mice at P14. Scale bar: 200 µm. Images are representative of 6 repeats.

**Extended Data Fig. 3.**
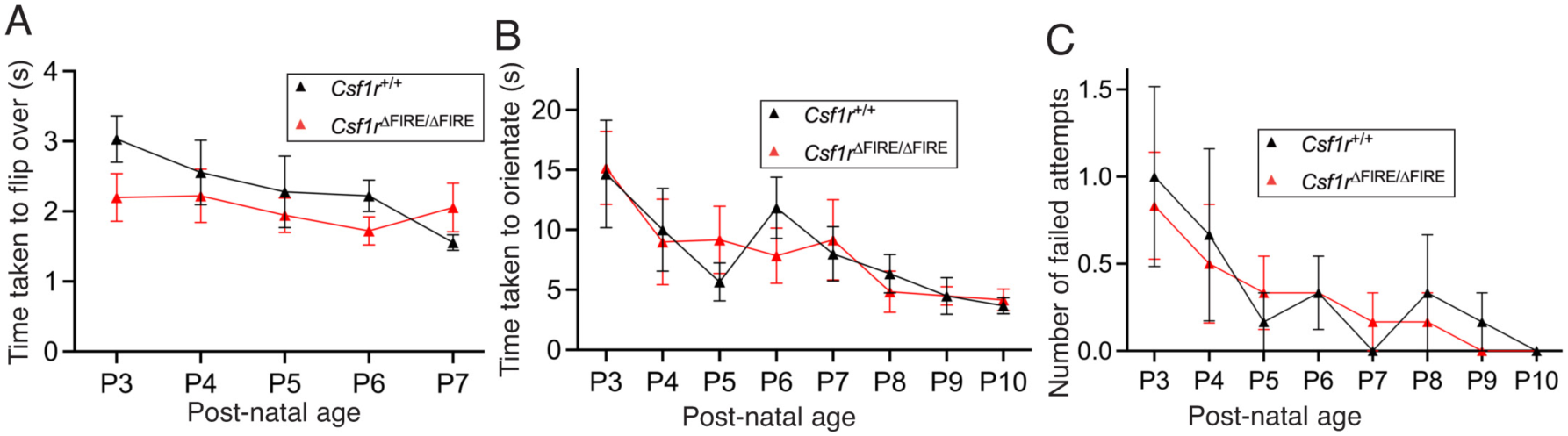
Assessing motor control in young pups. All data are shown as mean ± SEM. **A-C)** Mouse pup innate motor reflex testing. (A) shows time to turn over in righting reflex task. (B) and (C) relate to the negative geotaxis task, showing the time to reorientate uphill (B) and number of failed attempts (C); n=6 mice per genotype (3 males, 3 females).

**Extended Data Fig. 4.**
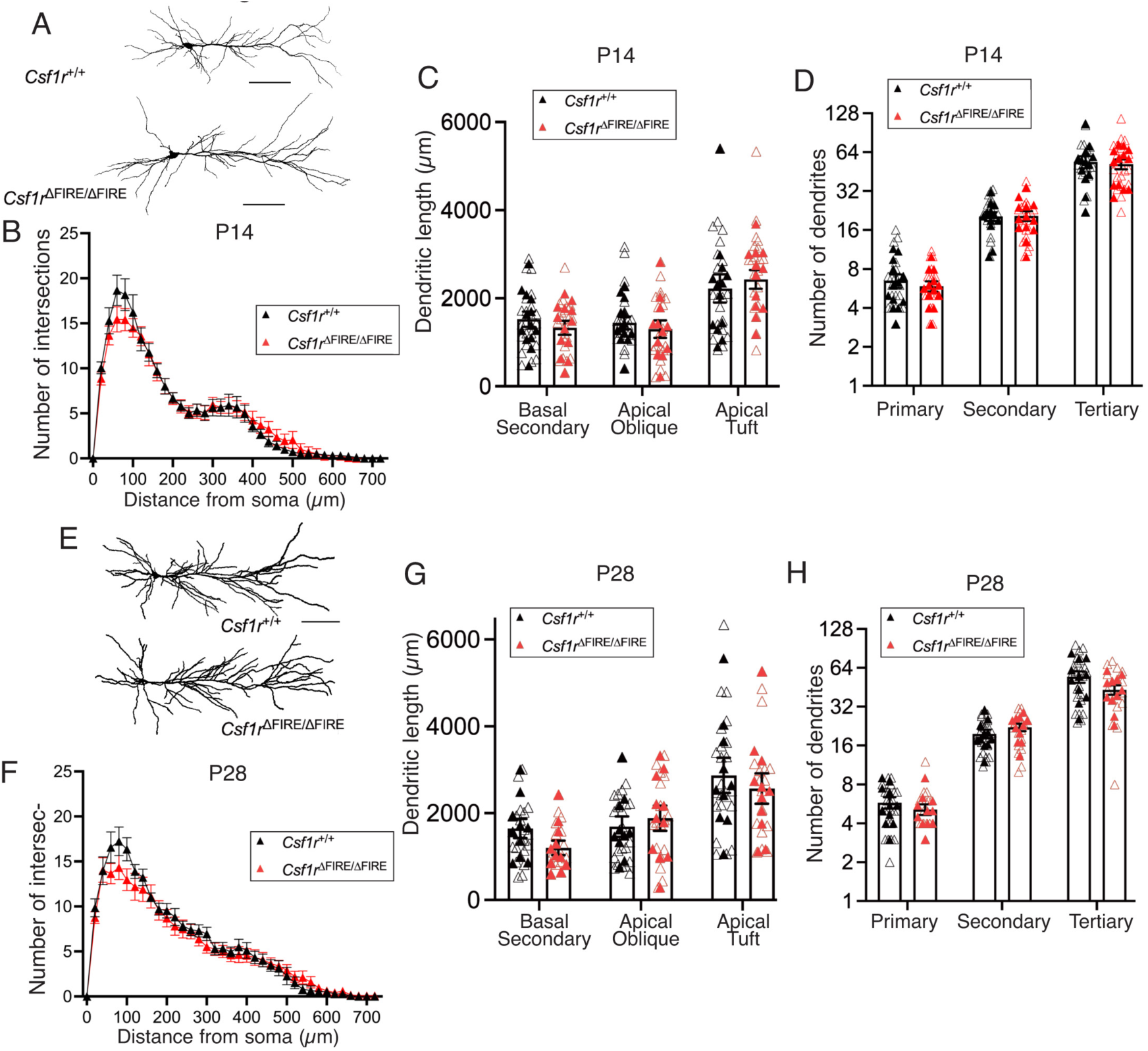
Absence of microglia during neurodevelopment does not lead to changes in CA1 pyramidal cell gross morphology. All data are shown as mean ± SEM and all statistical tests are two-sided. **A)** Example 2D reconstructions of the somato-dendritic axis of dye-filled CA1 pyramidal cells from *Csf1r*^+/+^(upper) and *Csf1r*^ΔFIRE/ΔFIRE^ (lower) P14 mice. Scale bars: 100 µm. **B**) Sholl analysis of reconstructed P14 CA1 pyramidal cell dendrites from *Csf1r*^+/+^ (n=13 mice) and *Csf1r*^ΔFIRE/ΔFIRE^ (n=13 mice). We observed no difference of Sholl distribution between *Csf1r*^+/+^ (black) and *Csf1r*^ΔFIRE/ΔFIRE^ (red) mice at P14 (F_(1,24)_=0.038, p=0.847, 2-way ANOVA [genotype])]; F_(36,864)_=0.826, p=0.758 [genotype_x_dendrite interaction]); n= 13 *Csf1r*^+/+^ (n=13 mice, 5m/8f) and 13 *Csf1r*^ΔFIRE/ΔFIRE^ mice (6m/7f). **C**) No difference in the length of difference dendritic types for reconstructed CA1 pyramidal cells at basal secondary, apical oblique, or apical tuft dendrites. Basal Secondary: F_(1,24)_=0.596, t_(24)_=0.772, p=0.448, nested Student’s t-test; Apical oblique: F_(1,24)_=1.161, t_(24)_=1.077, p=0.292, nested Student’s t-test; Apical tuft: F_(1,24)_=1.55, t_(24)_=1.245, p=0.225, nested Student’s t-test. **D**) No difference in number of dendrites for CA1 pyramidal cells (P14) was observed at primary, secondary or tertiary dendrites. Primary: F_(1,24)_=0.621, t_(24)_=0.788, p=0.439, nested Student’s t-test; Secondary: F_(1,24)_=0.143, t_(24)_=0.378, p=0.709, nested Student’s t-test; Tertiary: F_(1,24)_=0.020, t_(24)_=0.141, p=0.889, nested Student’s t-test. **E**) Reconstructions of CA1 pyramidal cells from P28 mice. **F**) No difference in Sholl distributions of CA1 pyramidal cell dendrites at P28 from *Csf1r*^+/+^ and *Csf1r*^ΔFIRE/ΔFIRE^ mice; F_(1,22)_=0.929, p=0.346, 2-way ANOVA [genotype]; F_(36,792)_=0.905, p=0.631 [genotype X distance interaction]. **G**) There was no change in the lengths of different dendrite types of CA1 pyramidal cells at P28 at basal secondary, apical oblique, or apical tuft dendrites. Basal secondary: F_(1,19)_=2.346, t_(19)_=1.532, p=0.142, nested Student’s t-test; Apical oblique: F_(1,19)_=0.350, t_(19)_=0.592, p=0.561, nested Student’s t-test; Apical tuft: F_(1,19)_=0.413, t_(19)_=0.643, p=0.528, nested Student’s t-test; n=10 *Csf1r*^+/+^ (4m/6f) and 11 *Csf1r*^ΔFIRE/ΔFIRE^ mice (6m/5f). **H**) We observed no difference in the number of primary secondary or tertiary dendrites at P28 in CA1 pyramidal cells. Primary: F_(1,20)_=0.172, t_(20)_=0.415, p=0.682, nested Student’s t-test; Secondary: F_(1,20)_=0.909, t_(20)_=0.954, p=0.352, nested Student’s t-test; Tertiary: F_(1,20)_=2.33, t_(20)_=1.53, p=0.143, nested Student’s t-test; n=11 *Csf1r*^+/+^ (5m/6f) and 11 *Csf1r*^ΔFIRE/ΔFIRE^ mice (6m/5f).

**Extended Data Fig. 5.**
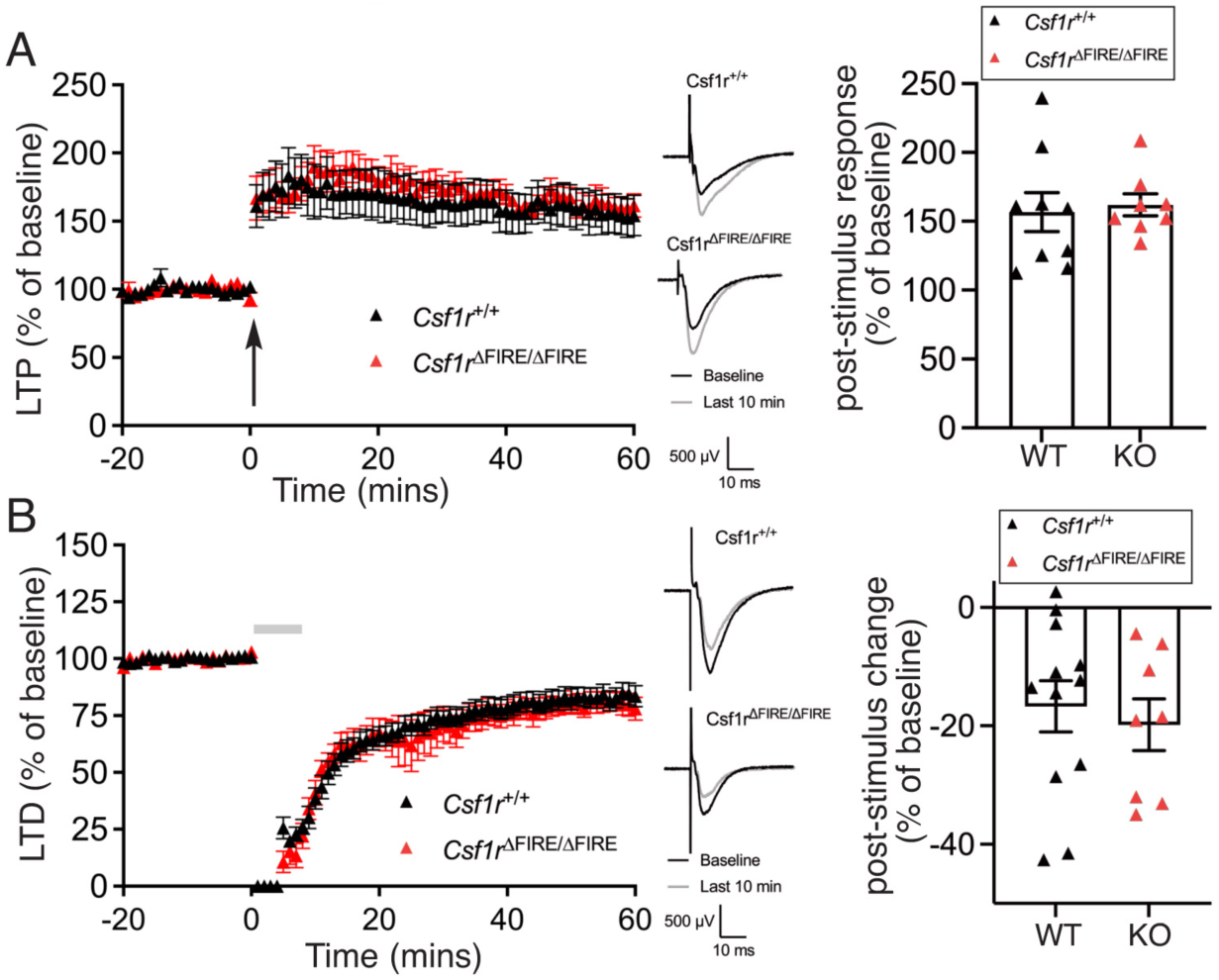
Analysis of synaptic plasticity. All data are shown as mean ± SEM and all statistical tests are two-sided. **A**) Time-course of long-term potentiation (LTP) at Schaffer-Collateral afferents in hippocampal area CA1, for *Csf1r*^+/+^ (black) and *Csf1r*^ΔFIRE/ΔFIRE^ (red) mice at P14. Data is shown for 1 hour following 2x 1 s 100 Hz tetanic stimulation (arrow). Example traces are shown for each genotype, before (black) and after (grey) LTP induction. Quantification of the magnitude of LTP for both genotypes revealed no difference (t_(15)_=0.31, p=0.761, 2-tailed Student’s t-test). N=9 *Csf1r*^+/+^ and 8 *Csf1r*^ΔFIRE/ΔFIRE^ mice. **B**) Time-course of long-term depression (LTD) of the Schaffer-Collateral afferents to CA1, induced by application of 50 µM R, S-DHPG (grey bar) in P14 mice. The magnitude of LTD, as measured 50-60 minutes after DHPG application revealed no effect of genotype (t_(18)_=0.486, p=0.633, 2-tailed Student’s t-test), n=12 *Csf1r*^+/+^ and 8 *Csf1r*^ΔFIRE/ΔFIRE^ mice.

**Extended Data Fig. 6.**
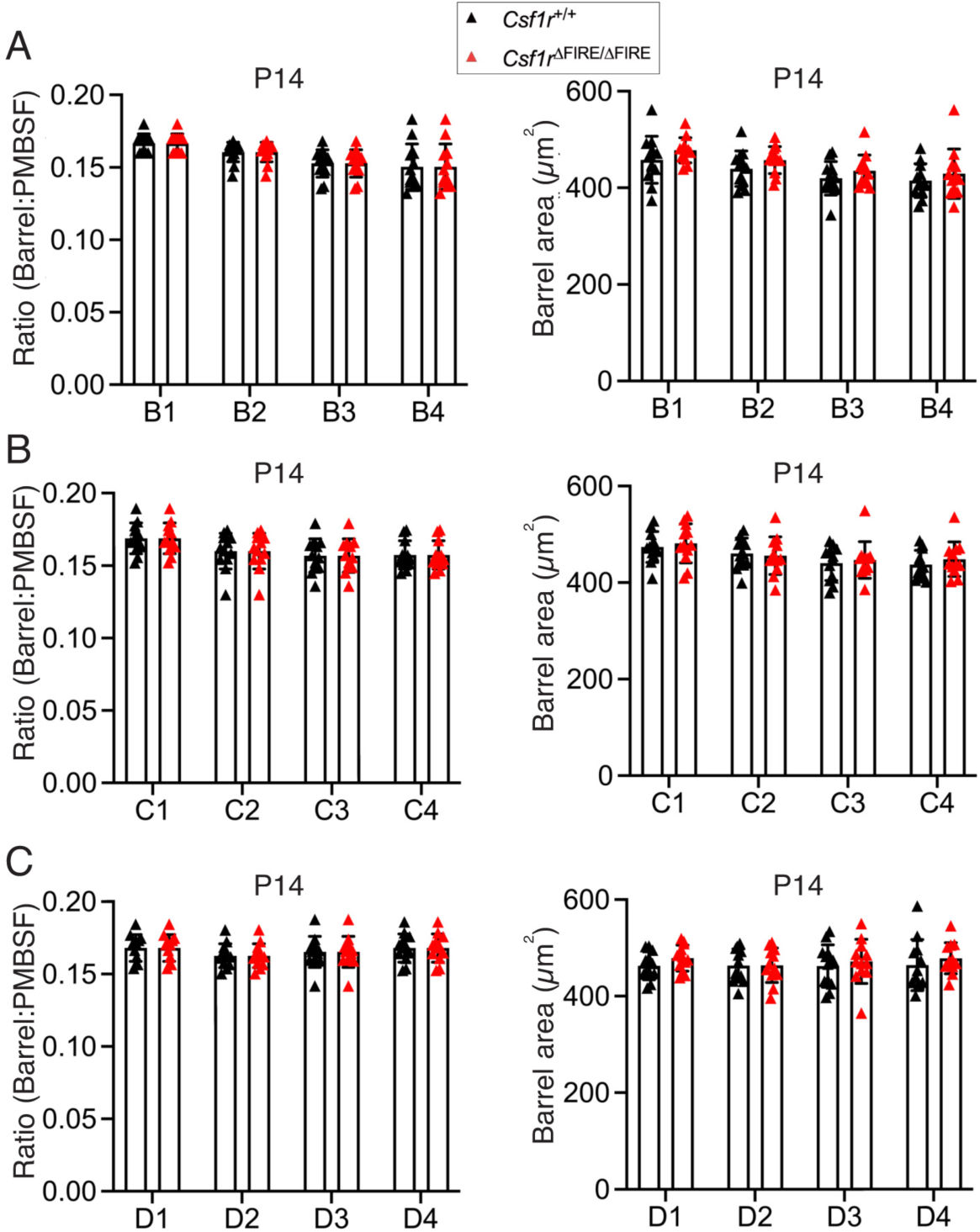
An absence of microglia does not alter the segregation of and size of individual somatosensory barrels. All data are shown as mean ± SEM and all statistical tests are two-sided. Ratio: The ratio of the barrel area in relation to the total area of the PMBSF. Barrel area: area of individual barrels 1-4 in rows B, C and D of the barrel field. **A-C)** Statistics (left-to right): F_(1, 22)_ = 3.26E-31, p>0.999; F_(1, 22)_ = 1.588, p=0.221 (A). F_(1, 22)_ = 1.63E-30, p>0.999; F _(1, 22)_ = 0.2076, p=0.653 (B). F_(1, 22)_ = 1.165E-29, p>0.999; F_(1, 22)_ = 0.557, p=0.464 (C). All are two-way ANOVAs [genotype effect], n=12 mice of each genotype.

**Extended Data Fig. 7.**
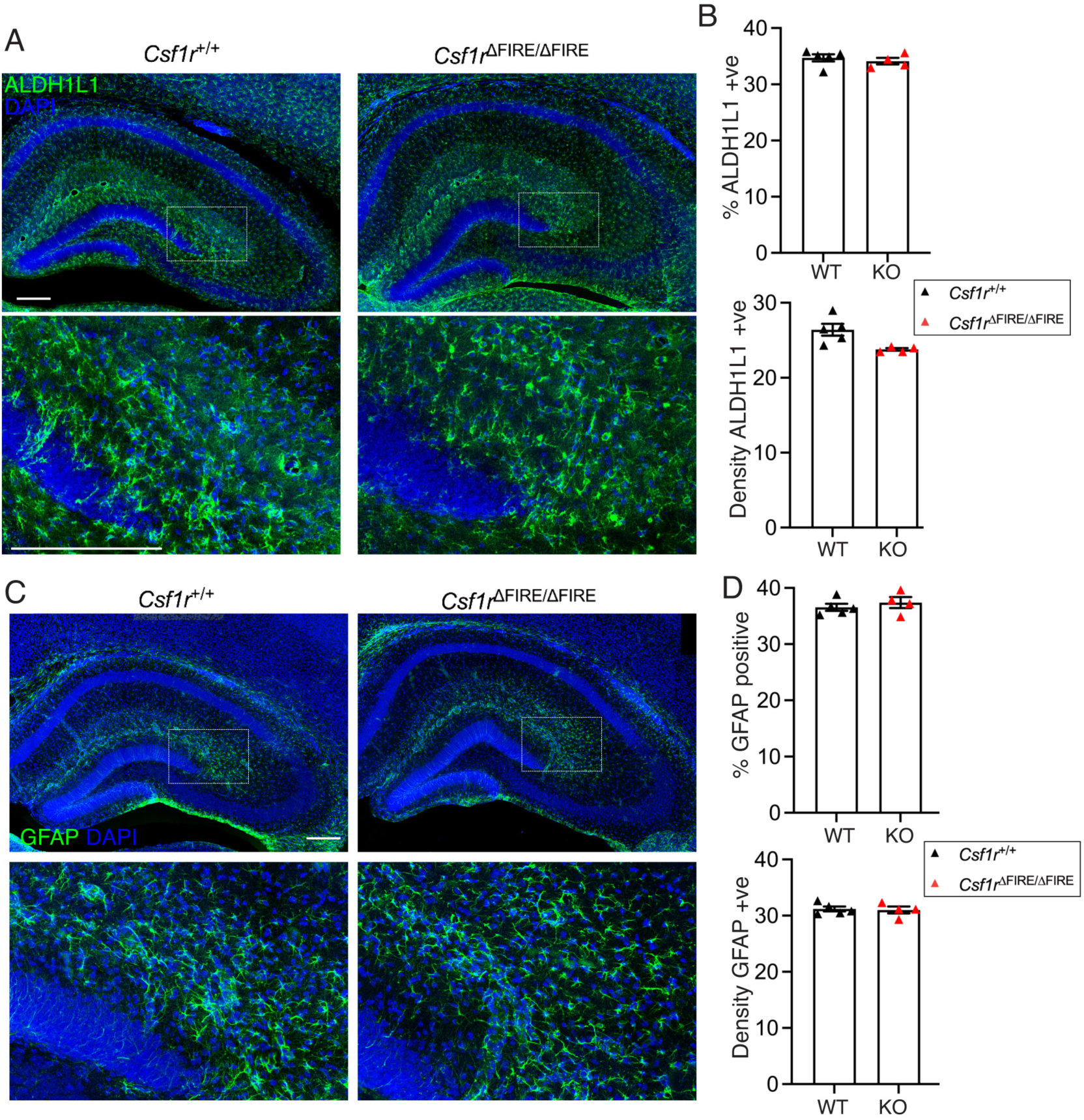
Analysis of astrocytes in the mouse hippocampus. All data are shown as mean ± SEM and all statistical tests are two-sided. **A, B)** Measurement of astrocyte density using an antibody against the pan-astrocyte marker ALDH1L1. Upper graph shows percent of DAPI-positive cells (t_(7)_=0.657, p=0.532 (unpaired t-test), lower graph shows density of astrocytes per 100 µm^2^ (t_(7)_=2.88, p=0.024 (unpaired t-test). Scale bar: 200 µm. **C, D)** Analysis as per (A, B) except that an antibody against GFAP was employed (t_(7)_=0.750, p=0.478 (upper); (t_(7)_=0.273, p=0.793 (lower, unpaired t-test), n=5 *Csf1r*^+/+^ (5m/2f) and n=4 (2m/2f) *Csf1r*^ΔFIRE/ΔFIRE^ mice. Scale bar: 200 µm.

**Extended Data Fig. 8.**
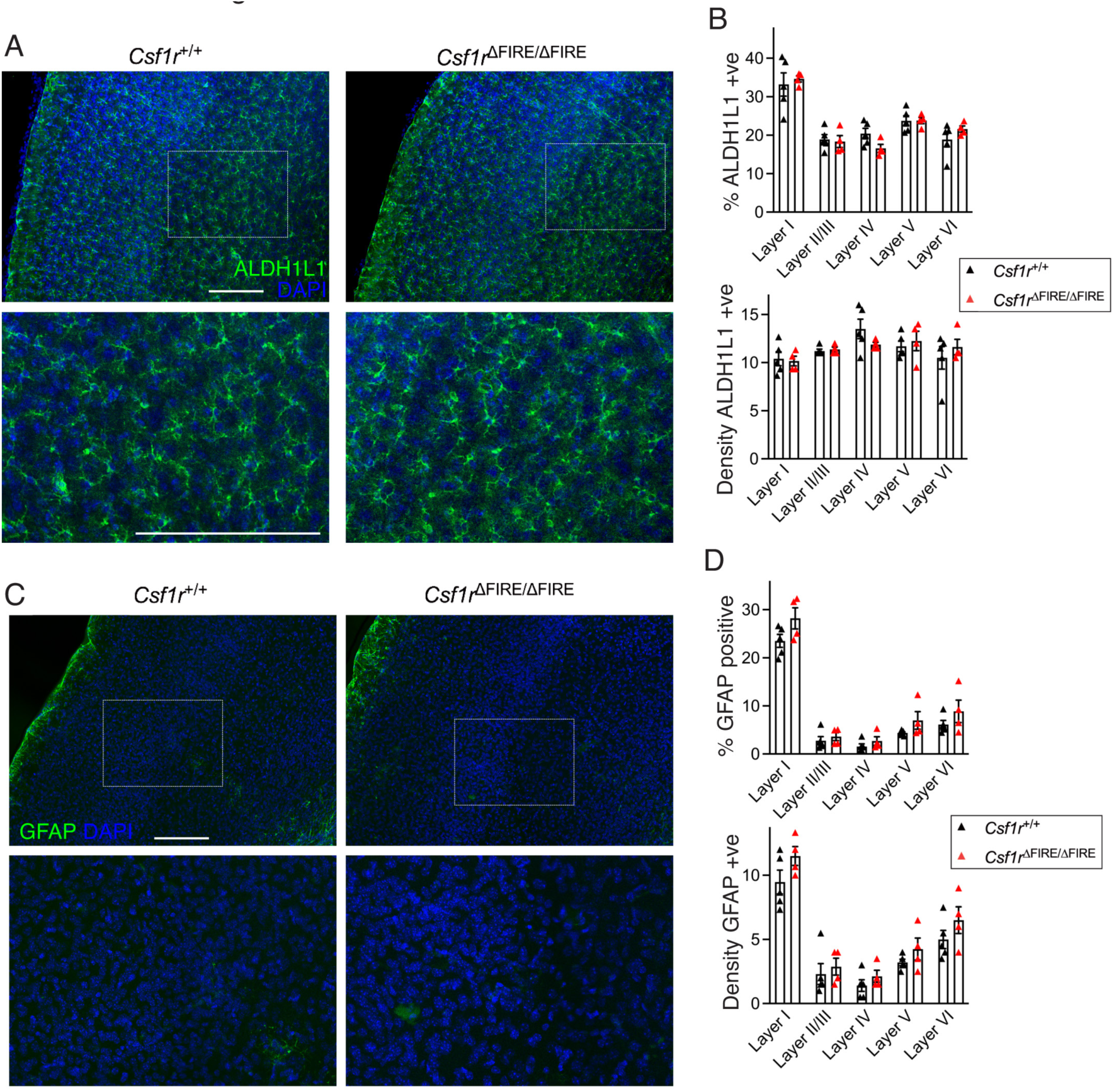
Analysis of astrocytes in the mouse neocortex. All data are shown as mean ± SEM and all statistical tests are two-sided. **A, B)** Measurement of astrocyte density using an antibody against the pan-astrocyte marker ALDH1L1. Upper graph shows percent of DAPI-positive cells; lower graph shows density of astrocytes (per 100 µm^2^). F_(1, 35)_=0.0009, p=0.977 (upper); F_(1, 35)_=0.1.2E-05, p=0.997 (lower, 2-way ANOVA, genotype effect). Scale bar: 200 µm **C, D)** Analysis as per (A, B) except that an antibody against GFAP was employed. C: F_(1, 35)_=8.81, p=0.0054. 2-way ANOVA, genotype effect). No individual layer showed statistical significance (Sidak’s post-hoc test). D: F_(1, 35)_=6.55, p=0.015, 2-way ANOVA, genotype effect). No individual layer showed statistical significance (Sidak’s post-hoc test). N=5 *Csf1r*^+/+^ (3m/2f) and n=4 (2m/2f) *Csf1r*^ΔFIRE/ΔFIRE^ mice). Scale bar: 200 µm.

## References

1 Sudhof, T. C. Towards an Understanding of Synapse Formation. Neuron 100, 276–293 (2018). 10.1016/j.neuron.2018.09.040

2 Hammond, T. R., Robinton, D. & Stevens, B. Microglia and the Brain: Complementary Partners in Development and Disease. Annu Rev Cell Dev Biol 34, 523–544 (2018). 10.1146/annurev-cellbio-100616-060509

3 Paolicelli, R. C. et al. Synaptic pruning by microglia is necessary for normal brain development. Science 333, 1456–1458 (2011). 10.1126/science.1202529

4 Hoshiko, M., Arnoux, I., Avignone, E., Yamamoto, N. & Audinat, E. Deficiency of the microglial receptor CX3CR1 impairs postnatal functional development of thalamocortical synapses in the barrel cortex. J Neurosci 32, 15106–15111 (2012). 10.1523/JNEUROSCI.1167-12.2012

5 Neniskyte, U. & Gross, C. T. Errant gardeners: glial-cell-dependent synaptic pruning and neurodevelopmental disorders. Nat Rev Neurosci 18, 658–670 (2017). 10.1038/nrn.2017.110

6 Weinhard, L. et al. Microglia remodel synapses by presynaptic trogocytosis and spine head filopodia induction. Nat Commun 9, 1228 (2018). 10.1038/s41467-018-03566-5

7 Stevens, B. et al. The classical complement cascade mediates CNS synapse elimination. Cell 131, 1164–1178 (2007). 10.1016/j.cell.2007.10.036

8 Schafer, D. P. et al. Microglia sculpt postnatal neural circuits in an activity and complement-dependent manner. Neuron 74, 691–705 (2012). 10.1016/j.neuron.2012.03.026

9 Wilton, D. K., Dissing-Olesen, L. & Stevens, B. Neuron-Glia Signaling in Synapse Elimination. Annu Rev Neurosci 42, 107–127 (2019). 10.1146/annurev-neuro-070918-050306

10 Favuzzi, E. et al. GABA-receptive microglia selectively sculpt developing inhibitory circuits. Cell 184, 5686 (2021). 10.1016/j.cell.2021.10.009

11 Cornell, J., Salinas, S., Huang, H. Y. & Zhou, M. Microglia regulation of synaptic plasticity and learning and memory. Neural Regen Res 17, 705–716 (2022). 10.4103/1673-5374.322423

12 Basilico, B. et al. Microglia shape presynaptic properties at developing glutamatergic synapses. Glia 67, 53–67 (2019). 10.1002/glia.23508

13 Filipello, F. et al. The Microglial Innate Immune Receptor TREM2 Is Required for Synapse Elimination and Normal Brain Connectivity. Immunity 48, 979–991 e978 (2018). 10.1016/j.immuni.2018.04.016

14 Lei, F. et al. CSF1R inhibition by a small-molecule inhibitor is not microglia specific; affecting hematopoiesis and the function of macrophages. Proc Natl Acad Sci U S A 117, 23336–23338 (2020). 10.1073/pnas.1922788117

15 Shah, R. et al. Metabolic Effects of CX3CR1 Deficiency in Diet-Induced Obese Mice. PLoS ONE 10, e0138317 (2015). 10.1371/journal.pone.0138317

16 Park, Y. et al. Fractalkine induces angiogenic potential in CX3CR1-expressing monocytes. Journal of leukocyte biology 103, 53–66 (2018). 10.1189/jlb.1A0117-002RR

17 Roy, C. et al. Shift in metabolic fuel in acylation-stimulating protein-deficient mice following a high-fat diet. Am J Physiol Endocrinol Metab 294, E1051–1059 (2008). 10.1152/ajpendo.00689.2007

18 Propson, N. E., Roy, E. R., Litvinchuk, A., Kohl, J. & Zheng, H. Endothelial C3a receptor mediates vascular inflammation and blood-brain barrier permeability during aging. J Clin Invest 131 (2021). 10.1172/JCI140966

19 Markiewski, M. M., Daugherity, E., Reese, B. & Karbowniczek, M. The Role of Complement in Angiogenesis. Antibodies (Basel) 9 (2020). 10.3390/antib9040067

20 Gorelik, A., Sapir, T., Woodruff, T. M. & Reiner, O. Serping1/C1 Inhibitor Affects Cortical Development in a Cell Autonomous and Non-cell Autonomous Manner. Front Cell Neurosci 11, 169 (2017). 10.3389/fncel.2017.00169

21 Gorelik, A. et al. Developmental activities of the complement pathway in migrating neurons. Nat Commun 8, 15096 (2017). 10.1038/ncomms15096

22 Hawksworth, O. A., Coulthard, L. G. & Woodruff, T. M. Complement in the fundamental processes of the cell. Mol Immunol 84, 17–25 (2017). 10.1016/j.molimm.2016.11.010

23 Rojo, R. et al. Deletion of a Csf1r enhancer selectively impacts CSF1R expression and development of tissue macrophage populations. Nat Commun 10, 3215 (2019). 10.1038/s41467-019-11053-8

24 Chitu, V. & Stanley, E. R. Regulation of Embryonic and Postnatal Development by the CSF-1 Receptor. Curr Top Dev Biol 123, 229–275 (2017). 10.1016/bs.ctdb.2016.10.004

25 Erblich, B., Zhu, L., Etgen, A. M., Dobrenis, K. & Pollard, J. W. Absence of colony stimulation factor-1 receptor results in loss of microglia, disrupted brain development and olfactory deficits. PLoS ONE 6, e26317 (2011). 10.1371/journal.pone.0026317

26 Godement, P., Salaun, J. & Imbert, M. Prenatal and postnatal development of retinogeniculate and retinocollicular projections in the mouse. The Journal of comparative neurology 230, 552–575 (1984). 10.1002/cne.902300406

27 Sretavan, D. & Shatz, C. J. Prenatal development of individual retinogeniculate axons during the period of segregation. Nature 308, 845–848 (1984). 10.1038/308845a0

28 Guido, W. Refinement of the retinogeniculate pathway. J Physiol 586, 4357–4362 (2008). 10.1113/jphysiol.2008.157115

29 Hong, Y. K. & Chen, C. Wiring and rewiring of the retinogeniculate synapse. Curr Opin Neurobiol 21, 228–237 (2011). 10.1016/j.conb.2011.02.007

30 Schecter, R. W. et al. Experience-Dependent Synaptic Plasticity in V1 Occurs without Microglial CX3CR1. J Neurosci 37, 10541–10553 (2017). 10.1523/JNEUROSCI.2679-16.2017

31 Chung, W. S. et al. Astrocytes mediate synapse elimination through MEGF10 and MERTK pathways. Nature 504, 394–400 (2013). nature12776 [pii] 10.1038/nature12776

32 Zhan, Y. et al. Deficient neuron-microglia signaling results in impaired functional brain connectivity and social behavior. Nat Neurosci 17, 400–406 (2014). 10.1038/nn.3641

33 Pagani, F. et al. Defective microglial development in the hippocampus of Cx3cr1 deficient mice. Front Cell Neurosci 9, 111 (2015). 10.3389/fncel.2015.00111

34 Mazaheri, F. et al. TREM2 deficiency impairs chemotaxis and microglial responses to neuronal injury. EMBO Rep 18, 1186–1198 (2017). 10.15252/embr.201743922

35 Lawrence, A. R. et al. Microglia maintain structural integrity during fetal brain morphogenesis. Cell 187, 962–980 e919 (2024). 10.1016/j.cell.2024.01.012

36 Gunner, G. et al. Sensory lesioning induces microglial synapse elimination via ADAM10 and fractalkine signaling. Nat Neurosci 22, 1075–1088 (2019). 10.1038/s41593-019-0419-y

37 Basilico, B. et al. Microglia control glutamatergic synapses in the adult mouse hippocampus. Glia 70, 173–195 (2022). 10.1002/glia.24101

38 Chu, Y. et al. Enhanced synaptic connectivity and epilepsy in C1q knockout mice. Proc Natl Acad Sci U S A 107, 7975–7980 (2010). 10.1073/pnas.0913449107

39 Surala, M. et al. Lifelong absence of microglia alters hippocampal glutamatergic networks but not synapse and spine density. EMBO Rep 25, 2348–2374 (2024). 10.1038/s44319-024-00130-9

40 Brown, T. C. et al. Microglia are dispensable for experience-dependent refinement of mouse visual circuitry. Nat Neurosci 27, 1462–1467 (2024). 10.1038/s41593-024-01706-3

## Methods references

41 Dobin, A. et al. STAR: ultrafast universal RNA-seq aligner. Bioinformatics 29, 15–21 (2013). bts635 [pii] 10.1093/bioinformatics/bts635

42 Liao, Y., Smyth, G. K. & Shi, W. featureCounts: an efficient general purpose program for assigning sequence reads to genomic features. Bioinformatics 30, 923–930 (2014). btt656 [pii] 10.1093/bioinformatics/btt656

43 Love, M. I., Huber, W. & Anders, S. Moderated estimation of fold change and dispersion for RNA-seq data with DESeq2. Genome Biol 15, 550 (2014). s13059-014-0550-8 [pii] 10.1186/s13059-014-0550-8

44 Zheng, G. X. et al. Massively parallel digital transcriptional profiling of single cells. Nat Commun 8, 14049 (2017). 10.1038/ncomms14049

45 Hao, Y. et al. Integrated analysis of multimodal single-cell data. Cell 184, 3573–3587 e3529 (2021). 10.1016/j.cell.2021.04.048

46 Young, M. D. & Behjati, S. SoupX removes ambient RNA contamination from droplet-based single-cell RNA sequencing data. Gigascience 9 (2020). 10.1093/gigascience/giaa151

47 Germain, P. L., Lun, A., Garcia Meixide, C., Macnair, W. & Robinson, M. D. Doublet identification in single-cell sequencing data using scDblFinder. F1000Res 10, 979 (2021). 10.12688/f1000research.73600.2

48 Thurman, A. L., Ratcliff, J. A., Chimenti, M. S. & Pezzulo, A. A. Differential gene expression analysis for multi-subject single-cell RNA-sequencing studies with aggregateBioVar. Bioinformatics 37, 3243–3251 (2021). 10.1093/bioinformatics/btab337

49 Oliveira, L. S., Sumera, A. & Booker, S. A. Repeated whole-cell patch-clamp recording from CA1 pyramidal cells in rodent hippocampal slices followed by axon initial segment labeling. STAR Protoc 2, 100336 (2021). 10.1016/j.xpro.2021.100336

50 Anderson, W. W. & Collingridge, G. L. Capabilities of the WinLTP data acquisition program extending beyond basic LTP experimental functions. J Neurosci Methods 162, 346–356 (2007). 10.1016/j.jneumeth.2006.12.018

51 Schindelin, J., et al. Fiji: an open-source platform for biological-image analysis. Nat Methods 9, 676–682 (2012). 10.1038/nmeth.2019

52 Longair, M. H., Baker, D. A. & Armstrong, J. D. Simple Neurite Tracer: open source software for reconstruction, visualization and analysis of neuronal processes. Bioinformatics 27, 2453–2454 (2011). 10.1093/bioinformatics/btr390

53 Sholl, D. A. Dendritic organization in the neurons of the visual and motor cortices of the cat. J Anat 87, 387–406 (1953).

54 Schikorski, T. & Stevens, C. F. Quantitative ultrastructural analysis of hippocampal excitatory synapses. J Neurosci 17, 5858–5867 (1997). 10.1523/JNEUROSCI.17-15-05858.1997

55 Csicsvari, J., Hirase, H., Czurko, A., Mamiya, A. & Buzsaki, G. Oscillatory coupling of hippocampal pyramidal cells and interneurons in the behaving Rat. J Neurosci 19, 274–287 (1999). 10.1523/JNEUROSCI.19-01-00274.1999

56 Zhang, X., Schlogl, A. & Jonas, P. Selective Routing of Spatial Information Flow from Input to Output in Hippocampal Granule Cells. Neuron 107, 1212–1225 e1217 (2020). 10.1016/j.neuron.2020.07.006

57 Malezieux, M., Kees, A. L. & Mulle, C. Theta Oscillations Coincide with Sustained Hyperpolarization in CA3 Pyramidal Cells, Underlying Decreased Firing. Cell Rep 32, 107868 (2020). 10.1016/j.celrep.2020.107868

58 Klausberger, T. et al. Brain-state– and cell-type-specific firing of hippocampal interneurons in vivo. Nature 421, 844–848 (2003). 10.1038/nature01374

59 Zeisel, A. et al. Molecular Architecture of the Mouse Nervous System. Cell 174, 999–1014 e1022 (2018). 10.1016/j.cell.2018.06.021

